# IRF1 regulates self-renewal and stress-responsiveness to support hematopoietic stem cell maintenance

**DOI:** 10.1101/2023.01.24.525321

**Authors:** Alexandra Rundberg Nilsson, Hongxu Xian, Shabnam Shalapour, Jörg Cammenga, Michael Karin

**Author notes:** Correspondence: Alexandra Rundberg Nilsson.

## Abstract

Inflammatory mediators induce emergency myelopoiesis and cycling of adult hematopoietic stem cells (HSCs) through incompletely understood mechanisms. To suppress the unwanted effects of inflammation and preserve its beneficial outcomes, the mechanisms by which inflammation affects hematopoiesis need to be fully elucidated. Rather than focusing on specific inflammatory stimuli, we here investigated the role of transcription factor Interferon (IFN) regulatory factor 1 (IRF1), which receives input from several inflammatory signaling pathways. We identify IRF1 as a master HSC regulator. IRF1 loss impairs HSC self-renewal, increases stress-induced cell cycle activation, and confers apoptosis resistance. Transcriptomic analysis revealed an aged, inflammatory signature devoid of IFN signaling with reduced megakaryocytic/erythroid priming and antigen presentation in IRF1-deficient HSCs. Finally, we conducted IRF1-based AML patient stratification to identify groups with distinct proliferative, survival and differentiation features, overlapping with our murine HSC results. Our findings position IRF1 as a pivotal regulator of HSC preservation and stress-induced responses.

## INTRODUCTION

Hematopoietic stem cells (HSCs) are the most hierarchically superior cells of the immune system. Their mostly quiescent nature, multi-lineage differentiation potential and self-renewal ability ensures lifelong production of blood cells^1^. Inflammatory signal transduction pathways play key roles in development, maintenance, and differentiation of HSCs. For instance, interleukin (IL)-1, as well as type I and type II interferons (IFNs), are involved in hematopoietic stem and progenitor cell (HSPC) specification during development^2^. Moreover, adult HSCs respond directly and indirectly to inflammatory cues, including tumor necrosis factor (TNF)^3–5^, IFNα^6^, IFNγ^7, 8^, polyinosinic:polycytidylic acid (pI:pC)^9^, lipopolysaccharide (LPS)^10^, and IL-1^11^ to induce mobilization, proliferation, and differentiation. This regulation is essential for maintenance of blood cell homeostasis and appropriate cellular responses to infections and injuries. The inability to support these functions can result in leukemia, aging-related phenotypes, an imbalanced blood system and chronic inflammation. Studies suggest that there may be a divergence in the HSC ability to recover from prolonged inflammatory challenge (by IL-1 and TNF) *versus* infectious stimuli (mimicked by LPS and pI:pC)^3, 9–11^. In-depth knowledge of inflammatory signaling is thus important to the understanding of both normal and abnormal HSC regulation, and for development of new therapeutic strategies aimed at inflammation-associated pathologies.

Interferon regulatory factor 1 (IRF1) is an evolutionally conserved transcription factor (TF), central to innate and adaptive immune responses^12^. IRF1 has mainly been studied in mature blood cells and hematopoietic organs and is presumed to exert its actions by controlling basal and inducible target gene expression. IRF1 was initially described for its involvement in IFNβ induction but has since been shown to be non-essential for type I IFN production^13, 14^. IRF1, however, has been linked to a plethora of other functions, including development, immune cell function, pattern recognition receptor (PRR) signaling, NLRP3 and AIM2 inflammasome activation, cell proliferation and growth, apoptosis, lipid metabolism, protein degradation, DNA damage and oncogenesis^15, 16^. IRF1 was also shown to control the transition from naive to formative pluripotency and susceptibility to viral infections in mouse embryonic stem cells^17^.

IRF1 expression is induced through several transcriptional mechanisms, including TNF signaling via IKK-NF-κB, RIG-I-like receptor (RLR) signaling via RIG-I-MAVS, Toll-like receptor (TLR) signaling via MYD88, and IFN signaling via JAK-STAT^15^. IRF1 binds together with other family members, such as IRF8, or other partners to IFN stimulated response elements (ISRE) or IRF-binding elements (IRF-E) at gene promoters to activate or repress transcription^18, 19^. The availability of IRF1 and its binding partners may differ between cell types in a stimulus-dependent manner, affecting its downstream consequences. Additionally, IRF1 can control gene expression through epigenetic alterations by interacting with histone-modification enzymes, such as the protein lysine acetylator p300^20^. IRF1 drives expression of IFN stimulated genes (ISGs) and cytokines, including antiviral, antibacterial, immune response^20–22^, apoptosis, and antigen presentation-related genes^16, 22^, and represses anti-proliferative genes. IRF1 thus represents a multifaceted protein at the center of inflammatory signaling and with the potential to control a wide array of cellular functions.

*Irf1* mRNA is expressed in most peripheral blood (PB) cells, including CD4^+^ and CD8^+^ T cells, NK cells, B cells and myeloid cells, and has confirmed roles in T cell development^23, 24^ and B cell expansion^25^. *Irf1* mRNA expression is also observed in HSCs (GSE14833 and GSE6506)^26^, although its role therein remains elusive. *Irf1* transcript levels increase during myeloid differentiation, and in response to inflammatory stimuli, including viral infections, LPS, type I and type II IFNs, IL-1, IL-12, and G-CSF^15, 27–34^. IRF1-deficient mice demonstrate decreased numbers of PB CD8^+^ T and NK cells, concomitant with increased numbers of CD4^+^ T cells^35, 36^. In the bone marrow (BM) compartment, *Irf1^-/-^* mice display increased numbers of immature granulocytic precursors with impaired granulocytic development^37^. Abnormal full-length *Irf1* transcript levels are associated with leukemia and myelodysplastic syndrome (MDS) with aberration of chromosome 5q (where the *Irf1* gene is located), breast cancer, and malignant melanoma^38^. The IRF1-responsive (Zhong et al., 2018) NLRP3 inflammasome has also been shown to function as a driver of MDS^39^ and mediate glucocorticoid resistance^40^. While reduced *Irf1* expression is linked to increased cell proliferation in leukemia, elevated *Irf1* upregulates programmed cell death ligand 1 (PD-L1) with subsequent evasion of immune surveillance in malignant melanoma^41^.

Heretofore, IRF1 has been investigated in several mature blood cell types but not in HSCs. Given the high relative expression of *Irf1* in HSCs compared to other BM progenitors, its multifaceted functions in mature blood cells, and its association with HSPC-originated leukemia and inflammation, we sought to define a role for IRF1 in HSC maintenance and regulation and identify the target genes through which it acts.

## RESULTS

### *Irf1^-/-^* mice show alterations in peripheral blood and bone marrow

We investigated the phenotypic effects of IRF1 ablation on the hematopoietic system in an unperturbed setting by comparing the composition of PB and BM cell types in primary *Irf1^-/-^* mice to wild-type (WT) controls (gating strategy is outlined in Supplementary Figure 1A-C). In accordance with previous reports^37^, we observed reduced CD8^+^ T and NK cell frequencies along with increased CD4^+^ T cell frequencies in *Irf1^-/-^* PB (Figure 1A). Furthermore, we observed an expanded myeloid fraction, primarily due to increased neutrophil abundance (Supplementary Figure 2A). In the BM, we found that HSC frequencies and LSKCD150^+^CD48^+^ (LKS^++^) cells were unaltered, while multipotent progenitors (MPPs) and granulocyte-monocyte lymphoid progenitors (GMLPs) were significantly decreased (Figure 1B). *Irf1*^-/-^ HSCs exhibited elevated CD150 expression (Figure 1C), a feature that is associated with myeloid-skewed, hierarchically-superior HSCs with increased long-term repopulation capacity, but also with functionally-declined, aged HSCs^42^. When investigating the frequencies of downstream intermediate and lineage-committed progenitors, we observed significantly reduced levels of common lymphoid progenitors (CLPs; Figure 1D). No differences were detected among pre-granulocyte-macrophage progenitors (PreGMs), granulocyte-macrophage progenitors (GMPs), pre-megakaryocytic-erythroid (PreMegE) progenitors, pre-colony-forming-unit-erythroid (PreCFU-E) precursors, megakaryocytic progenitors (MkPs), or colony-forming-unit-erythroid/proerythrocytes precursors (CFU-E/ProEry) (Figure 1D and Supplementary Figure 2B). Together, these results demonstrate significant deficiencies in the lymphoid compartments of both PB and BM and an altered HSC phenotype in *Irf1^-/-^* mice.

**Figure 1.**
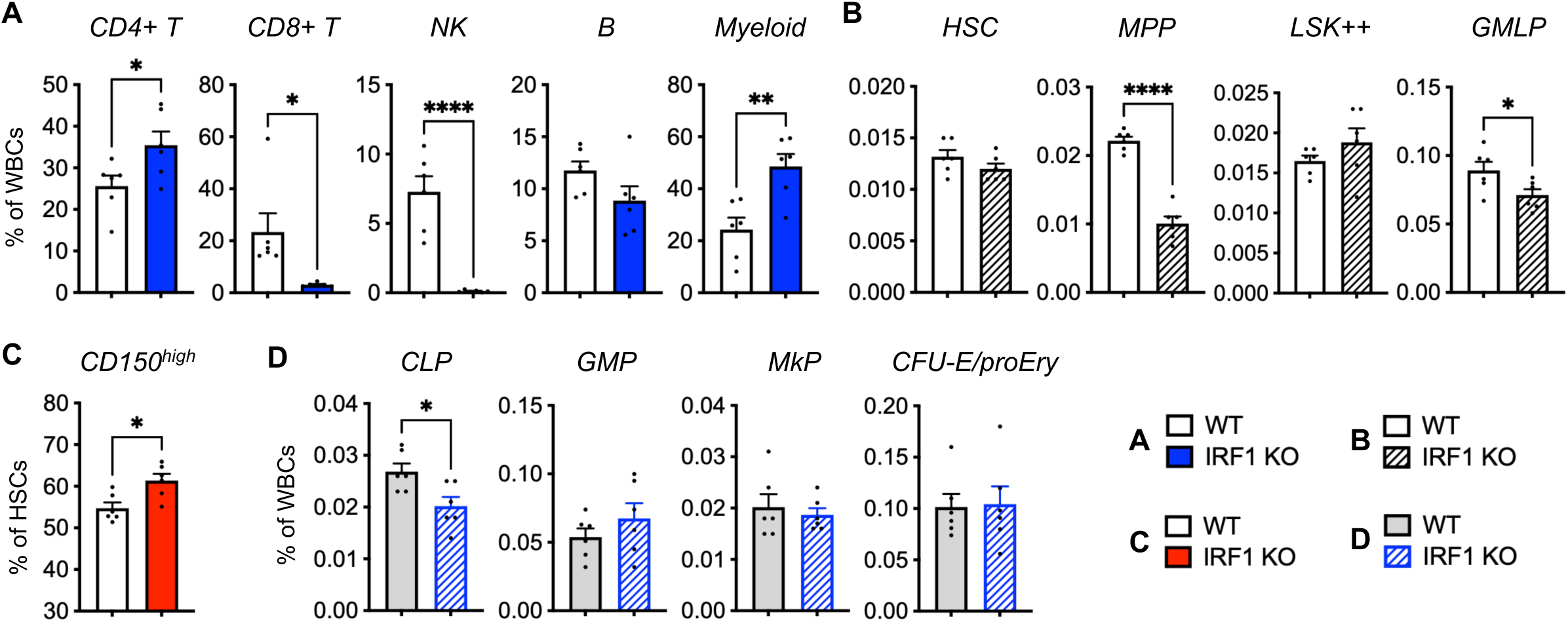
*Irf1*^-/-^ mice show alterations to the peripheral blood and bone marrow compartments. (A) Peripheral blood cell distribution within the CD45^+^ fraction. (B) Bone marrow LSK subpopulations frequencies within BM WBCs. (C) Fraction of CD150^high^ cells within the HSC population. (D) Fractions of lineage-committed progenitors within the bone marrow. WT n = 6, IRF1 KO n = 6. Error bars represent +SEM. *p < 0.05, **p < 0.01, ***p < 0.001, ****p < 0.0001. Littermate WT and *Irf1*^-/-^ mice were used for the experiment.

### *Irf1^-/-^* HSCs are more responsive to exogenous stress

We next evaluated the functionality of *Irf1^-/-^* HSCs. To this end, we assessed the HSC cell cycle phase distribution in primary steady-state mice, and observed no significant difference between WT and *Irf1^-/-^* HSCs (Figure 2A:b and 2B). We then subjected primary WT and *Irf1^-/-^* mice to exogenous inflammatory stress in the form of LPS, which activates IRF1 in macrophages^16, 43^, to evaluate inflammation-induced HSC activation. The mice were administered 35 μg of LPS by i.p. injections and sacrificed 16 hours later for HSC harvest and analysis (Figure 2A;c). *Irf1^-/-^* mice demonstrated significantly lower HSC activation/proliferation upon LPS exposure compared to WT mice (Figure 2C). We hypothesized that reduced HSC cell cycle activation in *Irf1^-/-^* mice could be indirect, i.e., due to diminished responses by HSC niche cells that produce HSC activating factors, rather than defective cell-autonomous LPS signaling. We therefore generated CD45.1-WT:CD45.2-*Irf1^-/-^* and CD45.1-WT:CD45.2-WT bone marrow chimeras to evaluate how WT and *Irf1^-/-^* HSCs responded to LPS within the same WT microenvironment. LPS administration was performed 12 weeks after transplantation (Figure 2A;d). Interestingly, in this setting, LPS-exposed CD45.2^+^ *Irf1^-/-^* HSCs exhibited significantly increased cell cycle activity compared to LPS-exposed CD45.2^+^ WT HSCs (Figure 2D). Importantly, no differences were observed between vehicle (PBS)-exposed CD45.2^+^ WT and *Irf1^-/-^* HSCs (Figure 2D), or between competitor CD45.1^+^ WT HSC populations within PBS- and LPS-treated groups, respectively (Supplementary Figure 3). Collectively, these results demonstrate that *Irf1^-/-^* HSCs are inherently more activatable by an inflammatory stimulus than WT HSCs.

**Figure 2.**
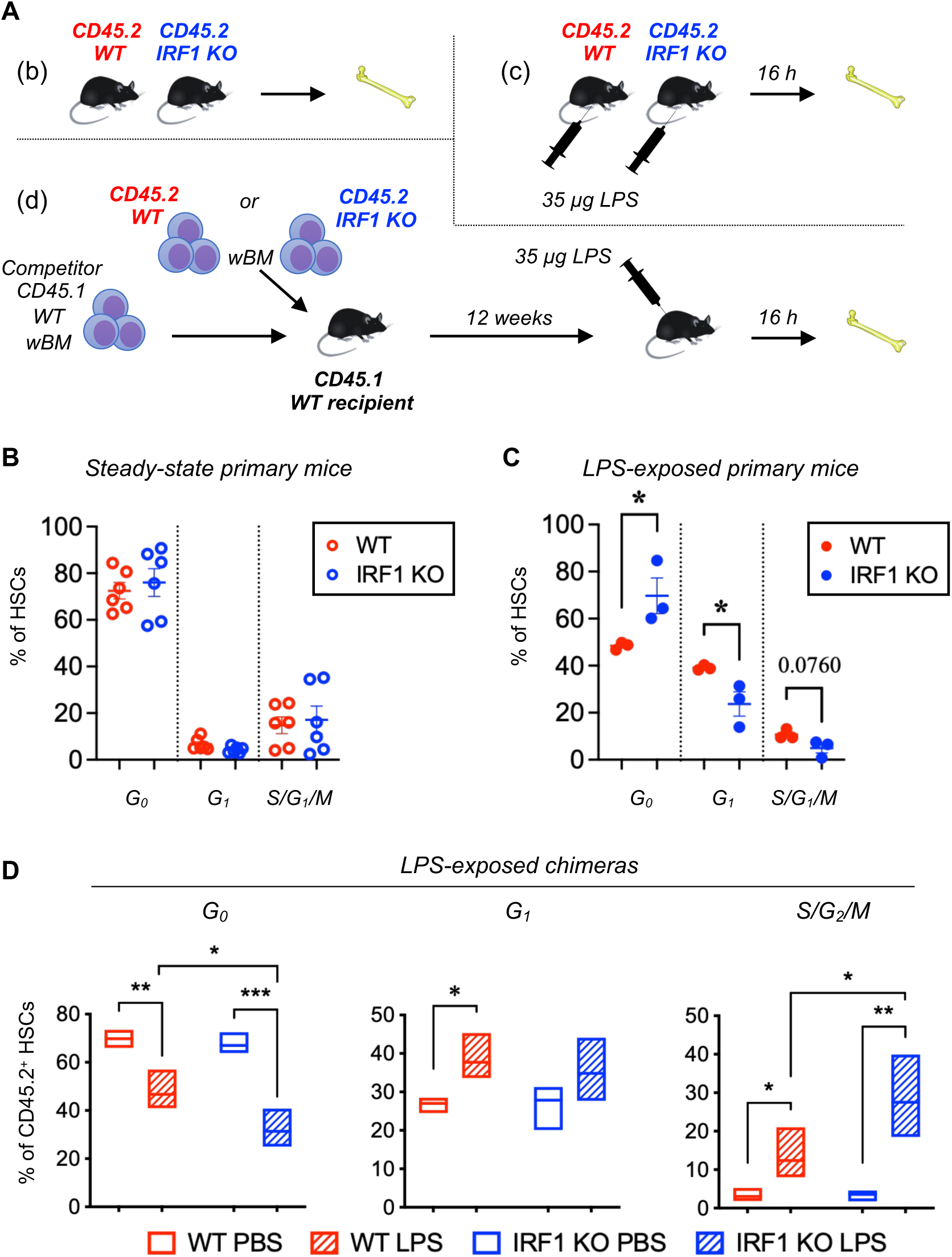
HSC cell cycle. (A) Experimental strategies for HSC cell cycle experiments. (B) HSC cell cycle distribution in steady-state WT and *Irf1*^-/-^ primary mice. WT n = 5, *Irf1*^-/-^ n = 5. (C) HSC cell cycle distribution in LPS-exposed WT and *Irf1*^-/-^ primary mice. WT n = 3, *Irf1*^-/-^ n = 3. Error bars represent ±SEM. D) Donor WT and *Irf1*^-/-^ CD45.2^+^ HSC cell cycle distribution in chimeric mice. WT PBS = 3, WT LPS = 5, *Irf1*^-/-^ PBS = 4, *Irf1*^-/-^ LPS = 3. *p < 0.05, **p < 0.01, ***p < 0.001, ****p < 0.0001. Box plots show floating bars (min to max) with line a mean. wBM = whole bone marrow.

### *Irf1^-/-^* HSCs show reduced long-term repopulation capacity

To further our investigations of altered *Irf1^-/-^* HSC functionality, we performed competitive HSC transplantations to evaluate the long-term self-renewal and reconstitution ability (Figure 3A). At early time points post transplantation, i.e. 4 and 8 weeks, WT and *Irf1^-/-^* HSCs gave rise to comparable chimerism levels in the PB CD4^+^ and CD8^+^ T cell and myeloid compartments (Figure 3B). *Irf1^-/-^* B cell chimerism was, however, considerably higher at all post-transplantation timepoints, which resulted in a prominent B cell biased distribution (Supplementary Figure 4A). This observation is likely linked to a previously reported increase in germinal center *Irf1*^-/-^ B cell expansion^25, 44^. A tendency towards reduced levels of myeloid cells could be observed beginning at 11 weeks post-transplantation (Figure 3B). A decline in myeloid cells that have a high turnover rate, but not of T cells which display much slower turnover, is in line with a pattern that can be observed upon HSC exhaustion. Indeed, when evaluating the chimerism within immature BM cells at endpoint, *Irf1^-/-^* chimerism levels were considerably reduced in all evaluated multipotent cell types, including HSCs (Figure 3C). Secondary competitive transplantation of re-purified HSCs confirmed a reduced long-term reconstitution capacity by *Irf1^-/-^* HSCs (Supplementary Figure 4B). These results suggest that exhaustion may account for the long-term repopulation deficiency of *Irf1^-/-^* HSCs.

**Figure 3.**
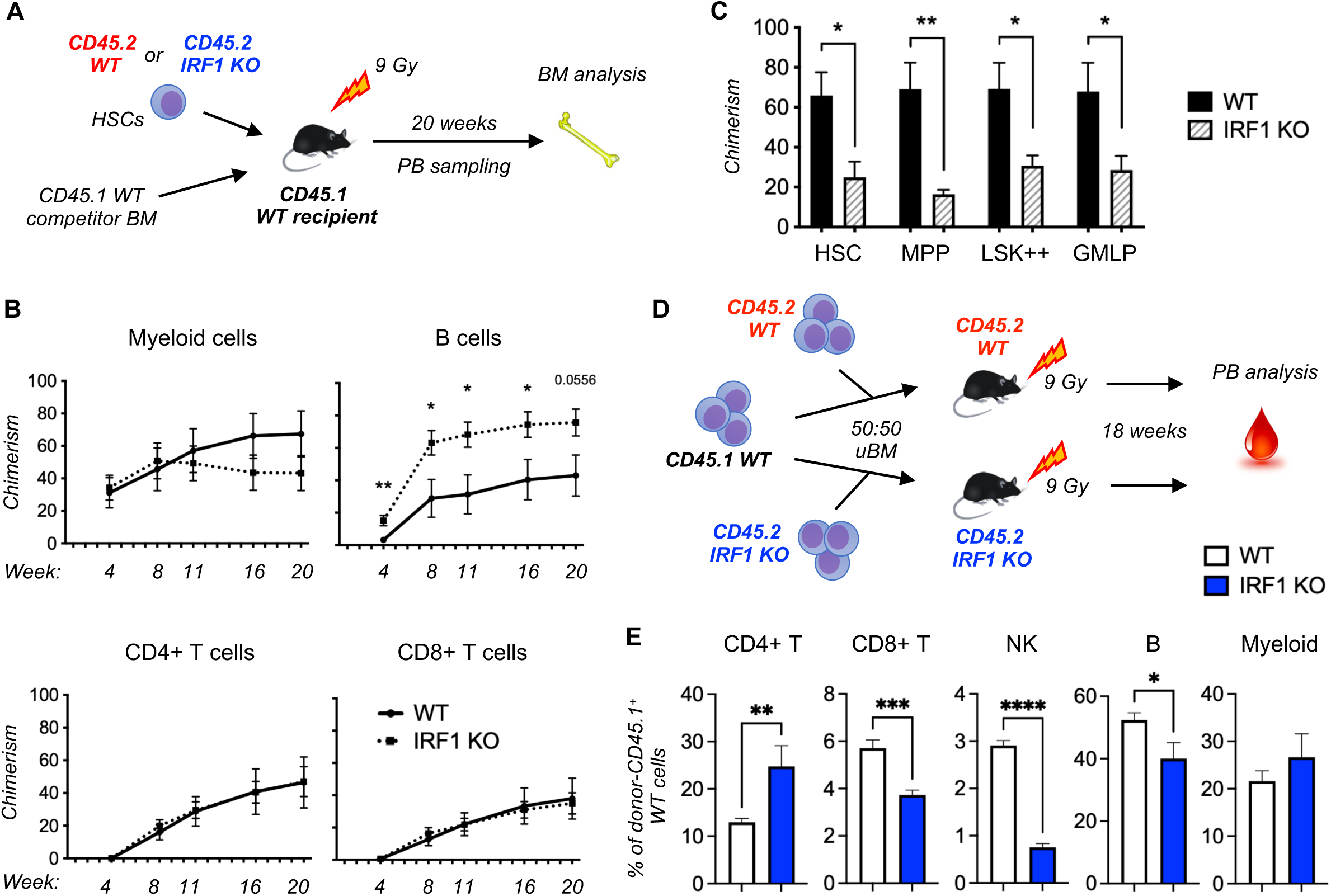
*Irf1*^-/-^ HSCs have reduced repopulation capacity. (A) Experimental strategy for competitive HSC transplantation experiment comparing WT and *Irf1*^-/-^ HSCs. Results are depicted in B and C. (B) CD45.2^+^ PB chimerism levels at various timepoints after competitive HSC transplantation. Error bars represent ±SEM. (C) Chimerism levels within the immature LSK subpopulation in the BM. Error bars represent +SEM. WT n = 6, *Irf1*^-/-^ n = 6. (D) Experimental strategy for the investigation of environmental impact on WT-derived blood cell output. Results are depicted in E. (E) CD45.1^+^ WT donor cell distribution within PB cell compartments. Error bars represent +SEM. Recipients: WT n = 8, *Irf1*^-/-^ n = 6. *p < 0.05, **p < 0.01, ***p < 0.001, ****p < 0.0001.

Because of differences in PB cell distribution between primary *Irf1^-/-^* mice (Figure 1A) and the *Irf1^-/-^* donor-derived output in transplanted mice (Supplementary Figure 4), we sought to evaluate the contribution from the *Irf1^-/-^* microenvironment to the observed PB perturbations. To this end, we transplanted CD45.1^+^ WT BM into CD45.2^+^ WT and *Irf1^-/-^* mice together with competing CD45.2^+^ WT and *Irf1^-/-^* BM, respectively (Figure 3D). CD45.1^+^ WT cells in the *Irf1^-/-^* microenvironment demonstrated vast alterations in PB lineage distribution compared to the same cells in the WT microenvironment (Figure 3E). These changes mimicked the alterations observed in primary *Irf1^-/-^* mice, including increased CD4^+^ T cell distribution, and reductions in CD8^+^ T and NK cells, suggesting that these alterations are extrinsically controlled by the *Irf1^-/-^* environment. Interestingly, the WT CD45.1^+^ B cell fraction was significantly reduced in the *Irf1^-/-^* environment (Figure 3E), which was not a consequence of competitive CD45.2^+^ *Irf1^-/-^* B cell expansion in the same mouse (data not shown). Together with the observations in primary mice (Figure 1A), these results the IRF1-deficient microenvironment inhibits lymphoid development.

### *Irf1^-/-^* HSCs show an altered inflammation-related transcriptomic profile

To uncover the transcriptomic underpinnings of altered *Irf1^-/-^* HSC functionality, we performed bulk RNA sequencing and analysis of purified WT and *Irf1^-/-^* HSCs (Figure 4A). Differentially expressed genes (DEGs) were identified using DESeq2 (adjusted p-value <0.05, log FC >0.58). With these criteria, we identified 169 upregulated and 134 downregulated genes in *Irf1^-/-^* HSCs (Figure 4B and C, Supplementary Tables 1 and 2). Ingenuity pathway analysis (IPA)^45^ predicted IRF1 together with the IRF1 direct targets STAT1 and TRIM24^22^ as the top 3 transcriptional regulators of the downregulated genes (Supplementary Table 3). In line with IRF1’s role in IFN signaling, downregulated genes were highly enriched for IFNα and IFNγ responses (Figure 4D, Supplementary Table 4). The downregulated genes were also enriched for apoptosis, antigen processing and presentation, and proteasome components (Figure 4D and E, Supplementary Tables 4 and 5). Conventional gene set enrichment analysis (GSEA) further revealed diminished unfolded protein response (UPR) and DNA repair in *Irf1^-/-^* HSCs (Supplementary Figure 5A).

**Figure 4.**
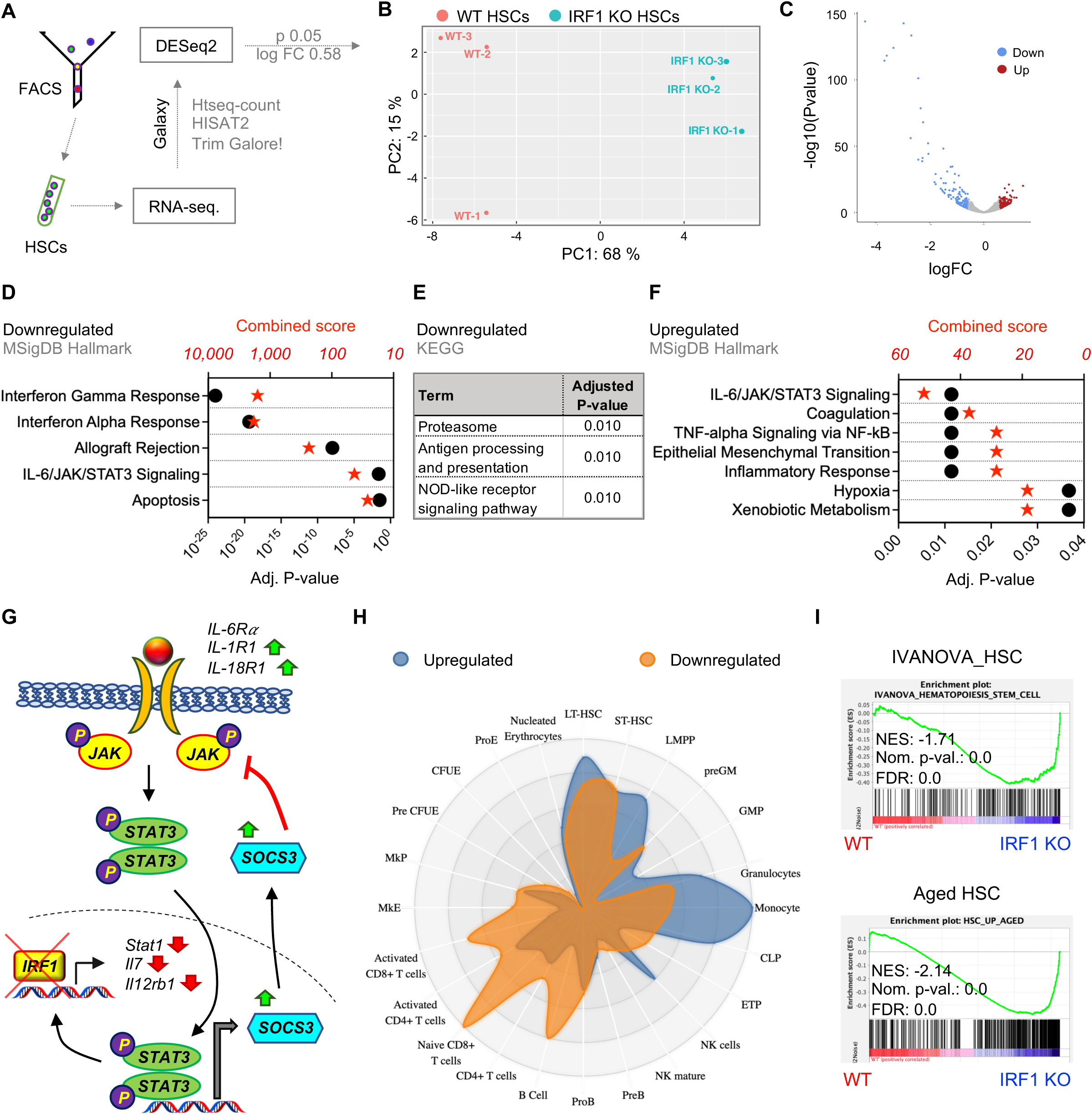
Gene expression analysis. (A) Experimental strategy for gene expression analysis. (B) PCA plot showing the relationship between the samples. (C) Volcano plot indicating differentially expressed genes (DEGs) between WT and *Irf1*^-/-^ HSCs. Red and blue dots represent significant DEGs with log FC > 0.58 and significance threshold <0.05. Enrichment of (D) MSigDB Hallmark and (E) KEGG gene sets within downregulated genes. (F) Enrichment of MSigDB Hallmark gene sets within upregulated genes. (G) Central Up- (green arrows) and downregulated (red arrows) genes in the IL-6/JAK/STAT2 pathway in *Irf1*^-/-^ HSCs. (H) Conventional GSEA on WT and *Irf1*^-/-^ gene expression profiles for lineage- and HSC-associated gene sets.

Interestingly, both up- and down-regulated genes were enriched for IL-6/JAK/STAT3 signaling (Figure 4D and E, Supplementary Tables 4 and 6). When investigating this gene set in detail, we found that most of the downregulated genes were located downstream to the IL- 6/JAK/STAT3 pathway, and had IRF1 binding sites^22^, whereas the upregulated genes did not have IRF1 binding sites and were situated upstream to the IL-6/JAK/STAT3 module or were direct STAT3 targets (Figure 4G). GSEA of WT *vs*. *Irf1^-/-^* HSCs revealed a significant overall enrichment of the IL-6/JAK/STAT3 pathway in *Irf1^-/-^* HSCs (Supplementary Figure 5B). These results highlight the complexity of signaling networks that mediate IRF1 function in HSCs.

Transcriptomic analysis revealed significant upregulation of additional inflammatory pathways in *Irf1^-/-^* HSCs, including TNF signaling. Interestingly, despite not observing significant differences in cell cycle status in primary *Irf1^-/-^* mice, the RNA-seq. data suggested a decreased proliferation-associated transcriptomic profile in *Irf1^-/-^* HSCs (Supplementary Figure 5C).

Given the increased CD150 expression on *Irf1^-/-^* HSCs and their perturbed lineage output, we next examined the enrichment of lineage-, HSC- and aging HSC-associated gene sets between WT and *Irf1^-/-^* HSCs. HSC CD150 expression was previously linked to aging, myeloid biased output, and megakaryocyte-primed gene expression^46^. Conversely, both lymphoid (CLP) and myeloid (PreGM) gene sets^47^ were enriched in *Irf1^-/-^* HSCs, while megakaryocytic (MkP) and erythroid (PreCFU-E) gene sets^47^ were downregulated (Supplementary Figure 5D, Supplementary Table 7). Cell type enrichment revealed that genes enriched for multipotent cell types, granulocytes, monocytes, and CLPs were upregulated, while genes associated with B and T cells were downregulated (Figure 4H). Interestingly, despite the observed reduction in HSC repopulation capacity that is frequently used as a surrogate measurement for stemness, the HSC gene set was also enriched in *Irf1^-/-^* HSCs (Figure 4I). However, a gene set characteristic of aged HSCs^47^, that are functionally inferior to young HSCs but also express higher levels of “stemness” genes, was also enriched in *Irf1^-/-^* HSCs (Figure 4I), whereas downregulated genes were enriched for HSC-associated genes (Figure 4H).

### *Irf1^-/-^* HSCs are less apoptotic, and express less ubiquitinated proteins and MHC class II

Reduced apoptosis of *Irf1^-/-^* HSCs, as suggested by the RNA-seq analysis, could potentially compensate for an impaired functionality, and contribute to the maintenance of HSCs in primary *Irf1^-/-^* mice and at early timepoints after transplantation. We therefore aimed to functionally confirm this observation by estimating the number of apoptotic cells by measuring annexin V^+^ cells by flow cytometry. Corroborating the transcriptional analysis, we detected reduced annexin V levels in *Irf1^-/-^* HSCs (Figure 5A and B). We then treated the cells with camptothecin before analysis and found that *Irf1^-/-^* HSCs exhibited lower levels of apoptosis compared to WT HSCs also in a setting of increased stress. These results show that *Irf1^-/-^* HSCs had acquired apoptosis resistance because of IRF1 ablation.

**Figure 5.**
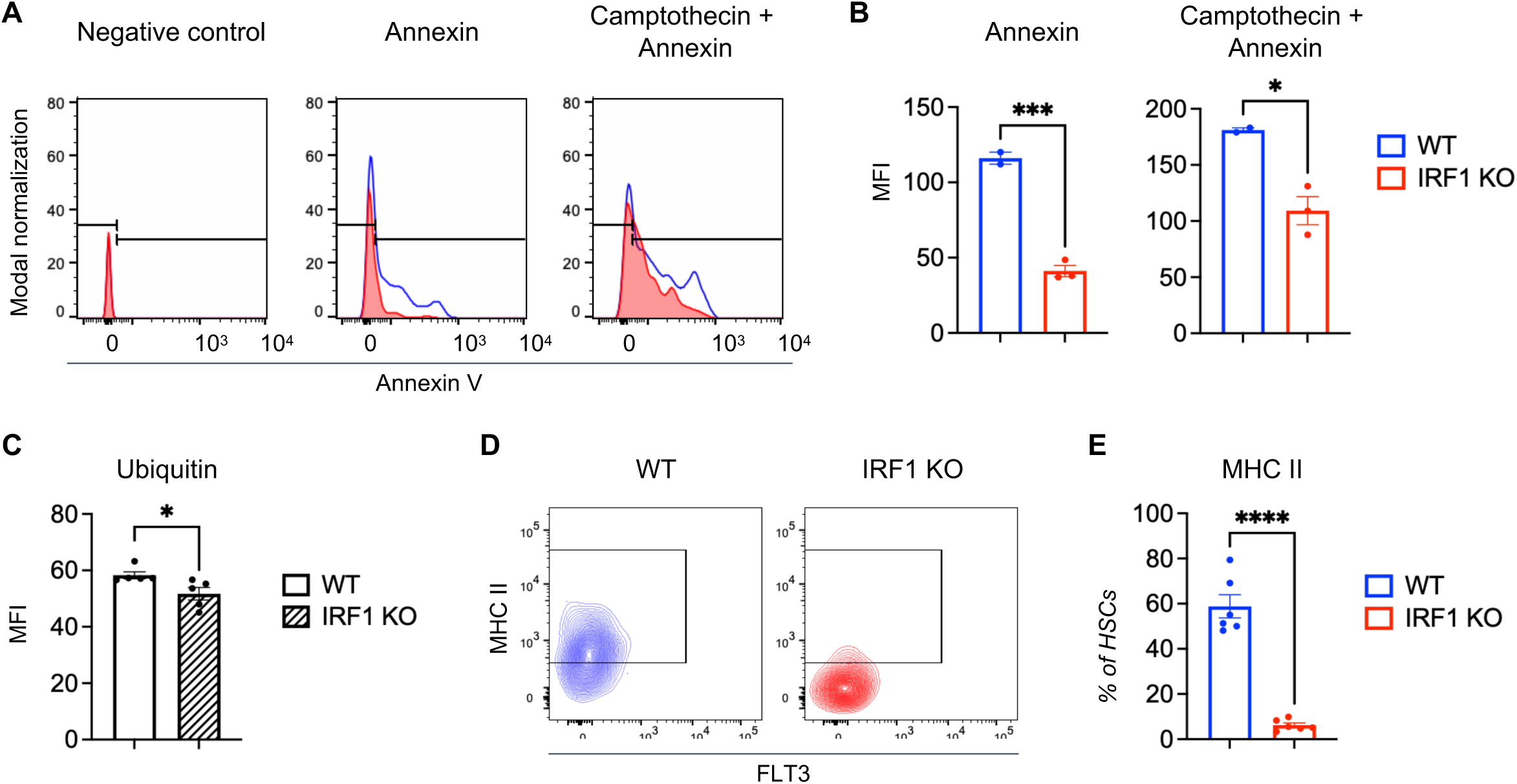
*Irf1*^-/-^ HSCs show improved survival, increased protein ubiquitylation, and decreased cell surface MHC class II expression. (A) Representative histograms Annexin V for WT (blue) and *Irf1*^-/-^ HSCs upon addition of Annexin and camptothecin. (B) Quantification of A. (C) Quantification of ubiquitin in WT and *Irf1*^-/-^ HSCs. (D) Representative plots visualizing the expression of MHC class II in WT and *Irf1*^-/-^ HSCs. (E) Quantification of the frequency of MHC II-positive HSCs in WT and *Irf1*^-/-^ mice. Error bars represent ±SEM. *p < 0.05, **p < 0.01, ***p < 0.001, ****p < 0.0001.

In further support of an altered functional state in *Irf1^-/-^* HSCs, the transcriptional analysis revealed reductions in UPR signaling, proteasome-associate transcripts and DNA repair. Dysregulations of these systems are linked to altered HSC function, transformation, and malignancy^48^. To validate reduced proteasome and UPR signaling, we measured the level of ubiquitinated proteins in WT and *Irf1^-/-^* HSCs. We found that *Irf1^-/-^* HSCs contained significantly less ubiquitin (Figure 5C, Supplementary Figure 5A).

Finally, our transcriptional profiling had demonstrated decreased expression of genes involved in antigen presentation and processing in *Irf1^-/-^* HSCs, including MHC class II genes *H2-Eb1, H2-Ab1, H2-Aa, H2-Q6, H2-DMa* and known controllers of MHC-II expression – *Ciita* and *Stat1* (Supplementary Table 1, Supplementary Figure 6). MHC-II expression was recently linked to HSC functionality^49, 50^, which prompted us to evaluate cell surface MHC-II expression on *Irf1^-/-^ HSCs*. We observed a striking 10.5-fold reduction in MHC-II on *Irf1^-/-^* HSCs (Figure 5D and E), suggesting that MHC-II may play a role in altered *Irf1^-/-^* HSC function.

### IRF1 expression marks distinct AML patient sample features

Given previous correlations between IRF1 dysregulation and myeloid malignancies^38, 51^, and our observation that IRF1-regulated HSC features are often hijacked by cancerous cells, we used a previously published set of 537 adult human AML samples (GSE5891) to conduct IRF1- based patient stratification. We allocated the AML samples into an IRF1^high^ group constituting the top 25 % expressing samples and a IRF1^low^ group composed of the 25 % bottom expressing samples with the remainder considered IRF1^medium^ (Supplementary Figure 7A). As expected based on the *IRF1* gene localization on chromosome 5, AMLs samples with loss of 5/7(q) were enriched in the IRF1^low^ group (Figure 6A). Moreover, IRF1^low^ AML were enriched in patients with t(8;21) and t(15;17) translocation, and contained fewer patients with a normal karyotype (NN). As observed in *Irf1^-/-^* HSCs, IRF1^low^ AML exhibited increased DNA repair/damage and survival, as well as increased proliferation concomitant with reduced differentiation and inflammatory signaling (Figure 6B and Supplementary Figure 7B). While further studies are required to properly decipher the role of IRF1 aberrations in leukemia, our results show that IRF1-based patient stratification marks functionally distinct AML subgroups which overlap with our *Irf1*^-/-^ HSCs findings.

**Figure 6.**
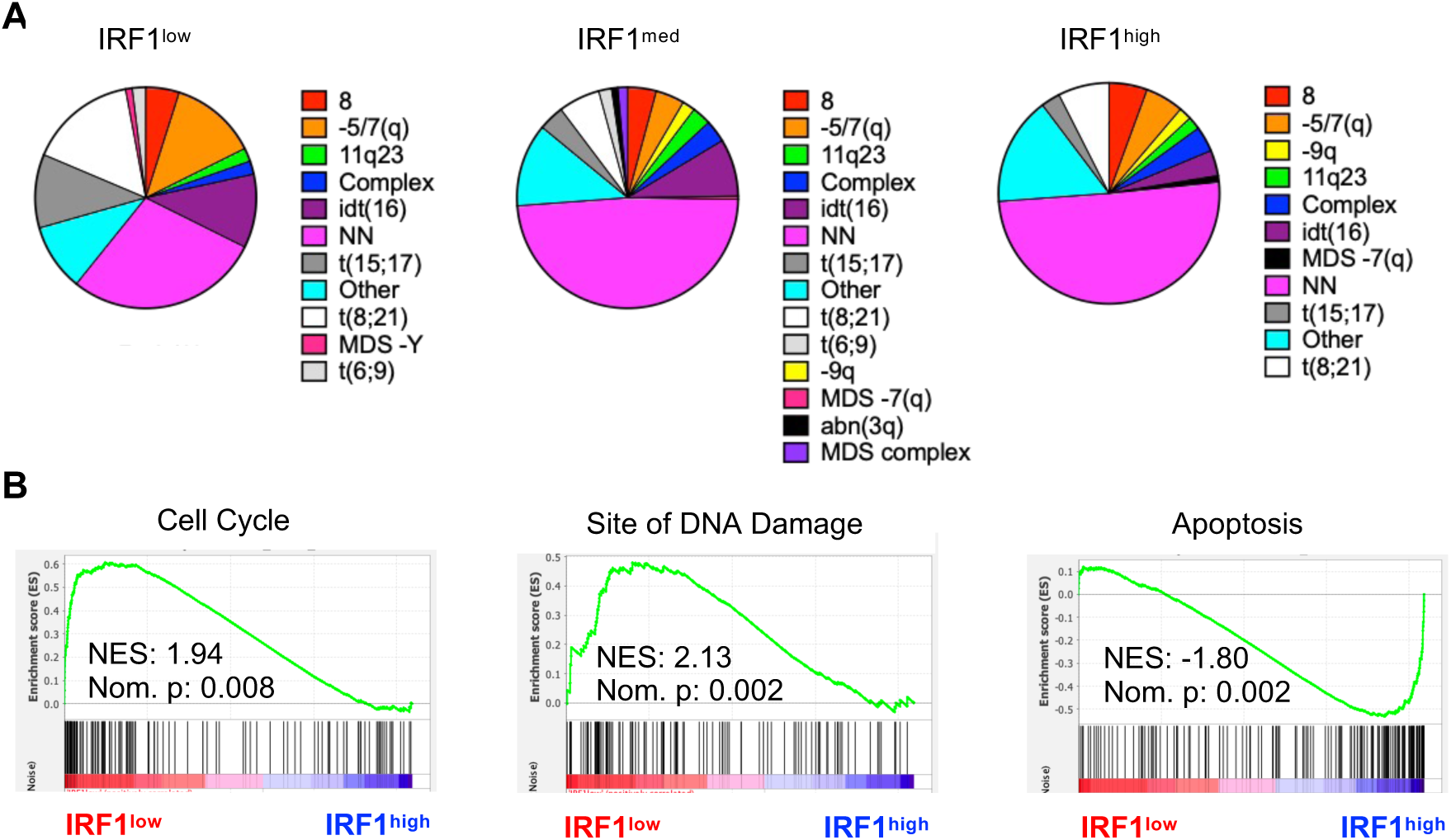
AML patient stratification based on IRF1 expression reveal distinct cancer- associated features. (A) Karyotype distribution within the three IRF1 expression level groups. 8 = trisomy 8, -5/7(q), -Y, -9q, -7(q) = loss of chromosomal segments, 11q23, t(15;17, t(8;21), t/6;9) = translocations, idt(16) = inversion, NN = normal karyotype, abn = abnormal, MDS = myelodysplastic syndrome. (B) GSEA between IRF1low and IRF1high patient samples for cell cycle, site of DNA damage, and apoptosis gene sets.

## DISCUSSION

IRF1 is associated with inflammation- and stress-triggered signaling pathways that promote IFN responses in mature blood cells, including macrophages. In this study, we report a novel role for IRF1 in the most primitive blood cells – the HSCs. We show that IRF1 is essential for long-term murine HSC maintenance and regulation of stress-related HSC responses (Figure 7). Although these aspects of IRF1 function was not previously investigated, IFNs and other inflammatory mediators that signal via IRF1-related pathways are known to have biological impact on HSCs. Reports of IRF1’s potential involvement in hematopoietic malignancies further hinted at a role in primitive hematopoietic compartments. IRF1 is, for instance, involved in NLRP3 inflammasome activation^16^, which drives MDS^39^. Additionally, IRF1 and other genes linked to inflammatory or intracellular stress signaling pathways that drive IFN production, such as DDX41 and SAMD9/L, are located on chromosomal segments that are commonly lost during myeloid malignant transformation^52–54^.

**Figure 7.**
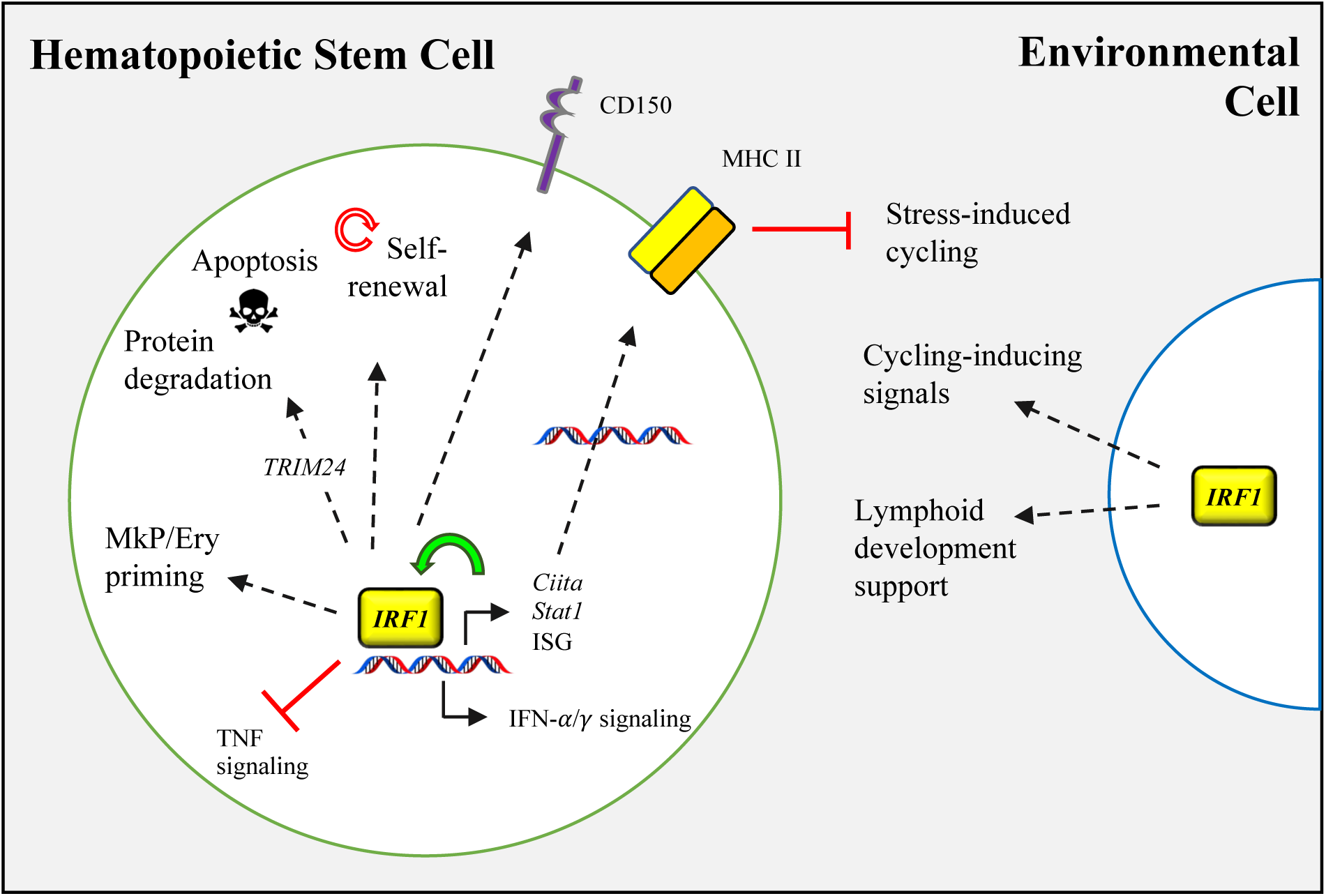
IRF1 controls fundamental HSC functions. IRF1 is a pivotal HSC regulator, controlling long-term self-renewal, apoptosis, protein degradation and inflammatory signaling. IRF1 transcriptionally primes HSCs towards the megakaryocytic and erythroid lineages, and regulates expression of key surface markers, including CD150 and MHC-II. Furthermore, IRF1 regulates lymphoid development and proliferation through extrinsic signaling.

We found that IRF1 is an essential regulator of apoptosis, DNA repair and protein degradation in HSC. Despite the reduced long-term maintenance of *Irf1^-/-^* HSCs, these cells displayed several features associated with elevated endurance, including reduced apoptosis, less DNA repair, and lower levels of ubiquitinated proteins and UPR. On the other hand, these findings may also be interpreted as an impaired ability to maintain cellular integrity and homeostasis, with consequential accumulation of cellular damage and reduced fitness. Nevertheless, these features may have partially compensated for an otherwise reduced long-term fitness, as observed in the unperturbed steady-state context and at early time-points post-transplantation. It is also plausible that loss of IRF1 can contribute to increased survival, clonal expansion, and mutagenesis in pre-leukemic cell clones if combined with leukemia-driving mutations.

IRF1-based AML patient stratification revealed distinct groups diverging in cancer-associated features, including proliferation, survival, DNA damage and lack of differentiation. Although further studies are required to demonstrate distinct treatment responses that are IRF1 dependent, such stratification may be applied to AMLs with different genomic aberrations and lead to discovery of more cost-efficient individualized treatment modalities.

It was demonstrated that MHC-II-mediated antigen presentation by HSCs to CD4^+^ T cells triggers their respective proliferation^49^. This interaction was further suggested to control differentiation and elimination of aberrant HSPCs. A later report revealed that MHC-II^low^ WT HSCs were more apoptotic and responsive to 5-FU- and pI:pC-induced proliferation^50^. We found that *Irf1^-/-^* HSCs exhibited reduced MHC-II expression and increased LPS-induced proliferation, suggesting that the latter feature may be controlled by an MHC-II-mediated mechanism. Moreover, loss of the direct IRF1 target STAT1^22^ in *Irf1*^-/-^ HSCs may account for the reduced MHC-II expression and impaired repopulation activity^50^. Loss of MHC-II promotes immune evasion^55^ resulting in decreased elimination of leukemic cells^49^.

We found that several of the PB perturbations associated with IRF1 loss, including reduced CD8^+^ T and NK cells, and increased CD4^+^ T cell numbers were not intrinsically primed but due to an extrinsic effect of the *Irf1^-/-^* microenvironment. Interestingly, the *Irf1^-/-^* microenvironment also suppressed B cell expansion, explaining why intrinsically primed *Irf1^-/-^* B cell expansion after transplantation into WT mice is not observed in primary *Irf1^-/-^* mice. The importance of separating direct *versus* indirect effects to delineate IRF1-dependent cellular features was also revealed by cell cycle analysis. While *Irf1^-/-^* HSCs in primary mice were impaired in their ability to cycle in response to LPS, experiments with WT: *Irf1^-/-^* chimeras revealed that this effect was due to reduced indirect responses by non-HSCs. *Irf1^-/-^* HSCs were in fact more responsive than WT HSCs when evaluated in a WT microenvironment. It is becoming increasingly clear that HSCs are regulated through different mechanisms than more mature blood cells, affecting their response to inflammatory stimuli^3, 4^. Deciphering inflammatory responses and the differences between various cell types is thus important to fully understand and find treatments for acute and chronic inflammatory and infectious conditions. This may be particularly crucial in the context of leukemia, where aberrant cells acquire stem cell characteristics that alter their modulation potential and therapeutic response.

In summary, the results presented in this study show that IRF1 is a key regulator of self-renewal, stress-responses, apoptosis, protein degradation, lineage priming, and inflammatory signaling in murine HSCs and as suggested by AML patient stratification these functions may extend to human AML cells (Figure 7).

## METHODS

### Mice

Age- and gender-matched *Irf1*^-/-^ mice (B6.129S2-Irf1^tm1Mak^/J), C57Bl/6, and B6.SJL- Ptprx^a^Pepc^b^/Boy mice were used throughout the study. The mice were maintained at the University of California San Diego (UCSD) Leichtag building vivarium. All procedures were conducted in accordance with ethical permit S00218 approved by Institutional Animal Care and Use Committee (IACUC).

### Isolation and analysis of peripheral blood and bone marrow

PB and BM were isolated and analyzed as previously described ^47, 56, 57^. Briefly, PB was collected in FACS buffer (PBS, 2 % FBS, 2 mM EDTA) supplemented with 0.0004 % heparin solution (STEMCELL Technologies Inc.). RBCs was removed by incubation at 37 °C for 25 minutes with 1 % Dextran solution in PBS, and room-temperature RBC lysis solution (STEMCELL Technologies Inc.). PB cells were stained with conjugated antibodies targeting CD4, CD8, NK1.1, CD19, CD11b, and Gr-1. BM cells were isolated from tibias, femurs and hip bones crushed with a pestle and mortar. Lineage depletion (B220, CD4, CD8, CD11b, Gr- 1, and -Ter119) or c-kit enrichment was performed using MACS magnetic microbead kit (Miltenyi Biotec) according to the manufacturer’s instructions. BM cells were stained with conjugated antibodies against B220, CD4, CD8, CD11b, Gr-1, Ter119, cKit, Sca1, CD48, CD150 APC, CD45.1, CD45.2, CD105 PE-Cy7, CD41, CD16/32, and MHC class II. Streptavidin-BV510 was used for biotin-identification. 1 μg/mL propidium Iodide (Molecular Probes) was used to distinguish viability. Flow cytometry analysis and assisted sorting was performed using Beckman Coulier (BC) CyAn ADP, Becton Dickinson (BD) LSR Fortessa X- 20, and BD Aria III.

### LPS treatments

Mice were administered 35 μg of LPS (Sigma Aldrich) i.p. and euthanized 16 hours after administration for downstream analysis. Control mice were administered 0.9 % saline solution.

### Transplantations

Competitive HSC transplantations were performed using 100 Lin^-^Sca-1^+^Kit^+^CD150^+^CD48^-^ HSCs together with 300 000 wBM competitor cells. wBM chimeras were generated with 10 million total donor BM cells. Recipient mice were lethally irradiated with 9 Gy and treated prophylactically with sulfamethoxazole and trimethoprim antibiotics for two weeks post transplantation in the drinking water. For serial transplantations, 276 donor-HSCs were re-isolated from each donor and transplanted together with 300 000 wBM competitor cells.

### Cell cycle analysis

Lineage-depleted BM cells were stained with HSC-identifying antibodies prior to fixation and permeabilization with BD Per/Wash^TM^ buffer (BD Pharmingen, Becton, Dickson and Company) at 4°C for 20 minutes. Cells were subsequently incubated with anti-Ki67 or IgG1 isotype control before wash and resuspension in 10 μg/mL 7-Aminoactinomycin D (7-AAD) diluted in Wash/Perm solution and incubated at 4°C over night before analysis.

### Apoptosis assay

BD Annexin V: FITC Detection kit I (BD Pharmingen) was used according to the manufacturer’s instructions. Briefly, lineage-depleted BM cells were stained with HSC-defining antibodies before washed and incubated with camptothecin at a final concentration of 5 μM for 5 hours at 37°C. The samples were then washed with ice-cold FACS buffer and resuspended in 1X Annexin binding buffer, to which Annexin V and PI were added. The samples were vortexed and incubated in the dark for 15 minutes at room temperature. Additional Annexin binding buffer was then added after which the samples were analyzed within one hour.

### Ubiquitin

After lineage depletion, antibody staining, and fixation, cells were incubated with anti-multi ubiquitin antibody for 30 minutes at 4°C. The cells were washed and resuspended with goat anti-mouse IgG Alexa Fluor 488 (Invitrogen) before analysis.

### RNA sequencing and analysis

∼1,600 HSC were sorted from each WT and IRF1 KO mouse into RLT lysis buffer supplemented with (1:100) beta-mercaptoethanol. RNA was isolated using the Single cell RNA purification kit (Norgen Biotek Corp.). cDNA generation and amplification were performed using the SMART-Seq® v4 Ultra® Low Input RNA Kit for sequencing (Takara Bio USA, Inc.). Quality check was performed using High sensitivity D5000 ScreenTape® on TapeStation Analysis Software 3.2 (Agilent Technologies, Inc.). Sequencing was performed using Nova Seq S4 (run type: PE100, type of library: NexteraXT, 25 million reads per sample; Illumina). Preprocessing and analysis were done using Galaxy Quality and adapter trimming of reads was performed using RNA Galaxy workbench 2.0^58^. Trim Galore! was used for quality check and adapter trimming of reads, HISAT2 for alignment and annotation, and Htseq-count to count aligned reads. Differentially expressed genes were identified using SESeq2. Volcano plots were generated with log FC = 0.58, significance threshold 0.05. The p value was adjusted for multiple testing with the Benjamini-Hochberg procedure which controls false discovery rate (FDR). Enrichment analysis of up- and downregulated genes was performed with Enrichr^59–61^. Global gene set enrichment analysis was performed with GSEA software^62, 63^. Upstream analysis was done with Qiagen IPA (https://digitalinsights.qiagen.com/IPA). Cell type enrichment was performed with CellRadar^64^.

### AML patient stratification

Expression data from GSE6891 containing 537 AML patient samples (<60 years of age) was used for patient stratification^65, 66^. 134 samples (∼25 %) with highest expression of IRF1 were allocated to the IRF1^high^ group, and 135 samples (∼25 %) with the lowest IRF1 expression to the IRF1^low^ group. The middle expressing samples (n = 268) were defined as IRF1^med^. Additional data about genetic aberrations were used to assess karyotype distribution between the 3 groups, which were further subjected to global gene set enrichment analysis.

### Quantification and statistical analysis

Statistical significance between experimental groups analyzed by flow cytometry was determined by unpaired student’s t-tests using Graphpad prism. Statistical details can be found in the figure legends. Division of mice into groups was randomized but not blind and no statistical methods were used to determine the number of mice for the experiments. Multi- lineage (T + B + My) donor reconstitution levels below 1 % in primary recipients were considered unsuccessful and excluded.

### Data sharing

RNA sequencing data is deposited under GEO accession number 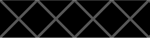. Further information and requests should be directed to the lead contact, Alexandra Rundberg Nilsson.

Detailed information about antibodies, reagents, and instruments can be found in Supplementary Table 8.

## ACKNOWLEDGEMENTS

We acknowledge lab manager Anne Poissant for her continuous support and essential work in the administrative managing of the laboratory. A.R.N. was supported by grants from the Tegger Foundation and the Swedish Society for Medical Research. M.K. was supported by grants from the NIH (AI043477).

## AUTHOR CONTRIBUTIONS

A.R.N designed and performed experiments, analyzed the data, and wrote the manuscript.

H.X., and S.S. helped with experiments and analyzes. M.K. supervised the study and wrote the manuscript. All co-authors provided input to the manuscript.

