## Supplementary files for "IRF1 regulates self-renewal and stress-responsiveness to support hematopoietic stem cell maintenance"

**SUPPLEMENTAL FIGURE AND TABLE LEGENDS**

**Supplementary Figure 1. Representative PB and BM gating strategies.** A) Representative gating strategy for PB. Pre-gated on singlets, viability, and scatter. Gating strategy for B) BM HSPCs and CLPs, and C) BM myeloid, erythroid and megakaryocytic precursors. Pre-gated on singlets, viability, scatter, and lineage negative.

**Supplementary Figure 2. *Irf1*$^{-/-}$ mice show alterations in hematopoietic compartments.** (A) Myeloid subpopulation frequencies within PB WBCs. (B) Intermediate progenitor population frequencies within BM WBCs. WT n = 6, *Irf1*$^{-/-}$ n = 6. Error bars represent +SEM. *p < 0.05, **p < 0.01, ***p < 0.001, ****p < 0.0001.

**Supplementary Figure 3. Partial presence of *Irf1*$^{-/-}$ hematopoietic cells does not alter LPS-induced proliferation responses by co-occurring WT HSCs.** (A) Endogenous and competitor WT donor CD45.1 HSC cell cycle distribution in chimeric mice. WT PBS = 3, WT LPS = 5, *Irf1*$^{-/-}$ PBS = 4, *Irf1*$^{-/-}$ LPS = 3. Exp#173. Error bars represent +SEM. *p < 0.05, **p < 0.01, ***p < 0.001, ****p < 0.0001.

**Supplementary Figure 4. *Irf1*$^{-/-}$ HSCs have a B-cell biased mature PB output and have impaired serial reconstitution capacity.** (A) CD45.2+ PB distribution levels at 20 weeks post competitive HSC transplantation. WT n = 6, *Irf1*$^{-/-}$ n = 6. (B) HSC chimerism levels at 23 weeks post serial HSC transplantation. WT n = 6, *Irf1*$^{-/-}$ n = 6. Error bars represent +SEM. *p < 0.05, **p < 0.01, ***p < 0.001, ****p < 0.0001.

**Supplementary Figure 5. Gene expression analysis.** GSEA of (A) unfolded protein response (UPR) and DNA repair, (B) IL6/JAK/STAT3 signaling and (C) proliferation-associated gene sets between WT and *Irf1*$^{-/-}$ HSCs.

**Supplementary Figure 6. *Irf1*$^{-/-}$ HSC show altered antigen presentation and processing.** Pink encircling indicates significant downregulation in *Irf1*$^{-/-}$ HSCs.

**Supplementary Figure 7.** (A) Patient stratification of the 537 AML patient samples based on IRF1 expression. (B) GSEA between IRF1$^{low}$ and IRF1$^{high}$ patient samples for DNA repair, regulation of leukocyte differentiation, and inflammatory response gene sets.

**Supplementary Table 1.** Significantly downregulated genes in *Irf1*$^{-/-}$ compared to WT HSCs.

**Supplementary Table 2.** Significantly upregulated genes in *Irf1*$^{-/-}$ compared to WT HSCs

**Supplementary Table 3.** Ingenuity Pathway Analysis (IPA) for significantly downregulated genes in *Irf1*$^{-/-}$ HSCs.

**Supplementary Table 4.** Significantly enriched MSigDB Hallmark pathways among downregulated genes identified with Enrichr.

**Supplementary Table 5.** Significantly enriched KEGG pathways among downregulated genes identified with Enrichr.

**Supplementary Table 6.** Significantly enriched MSigDB Halllmark pathways among upregulated genes identified with Enrichr.

**Supplementary Table 7.** Lineage- and HSC aging-associated gene sets used for GSEA.

**Supplementary Table 8.** Key resources table containing detailed information about antibodies, reagents, data and instruments used in the study.

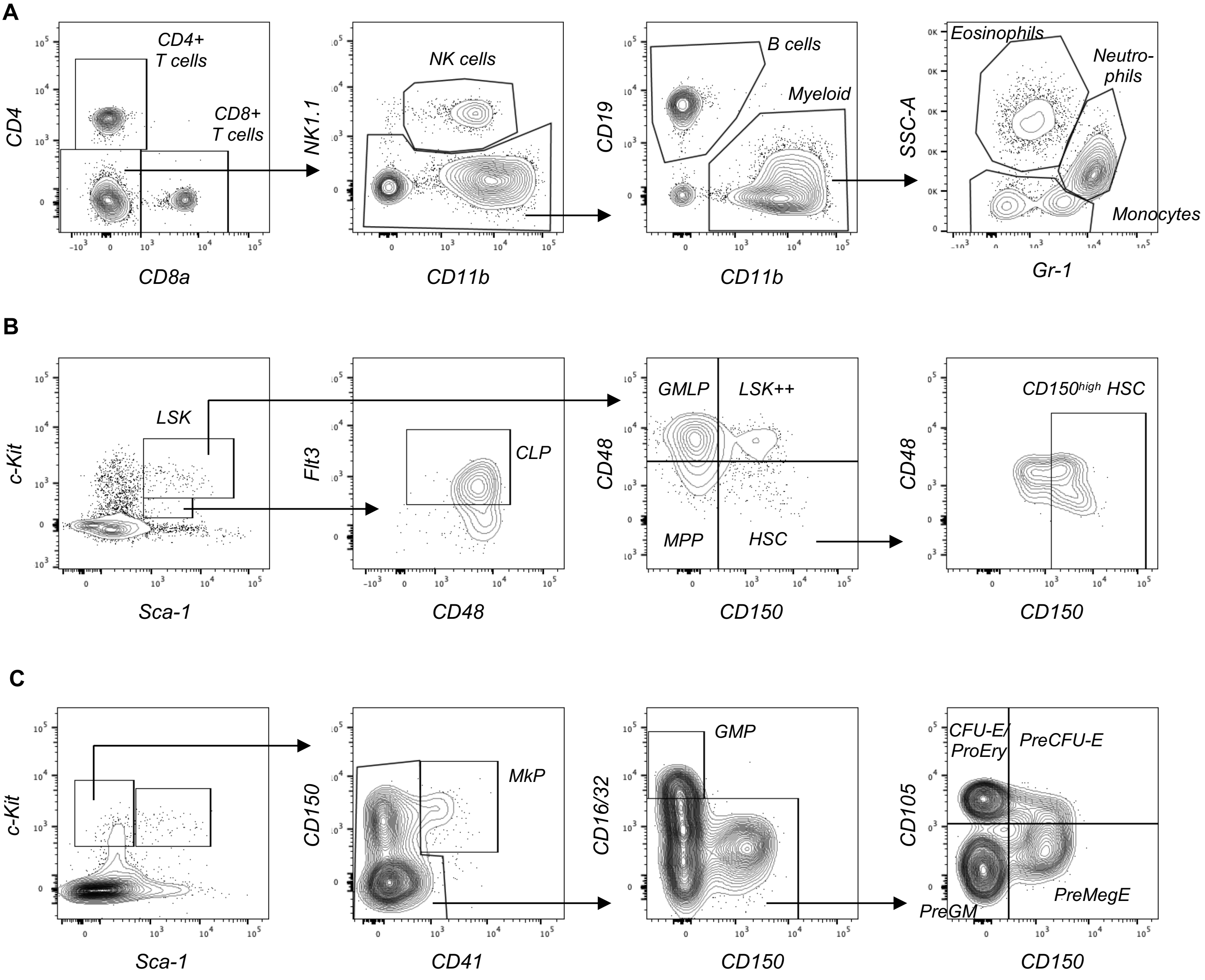

**Supplementary Figure 1.**

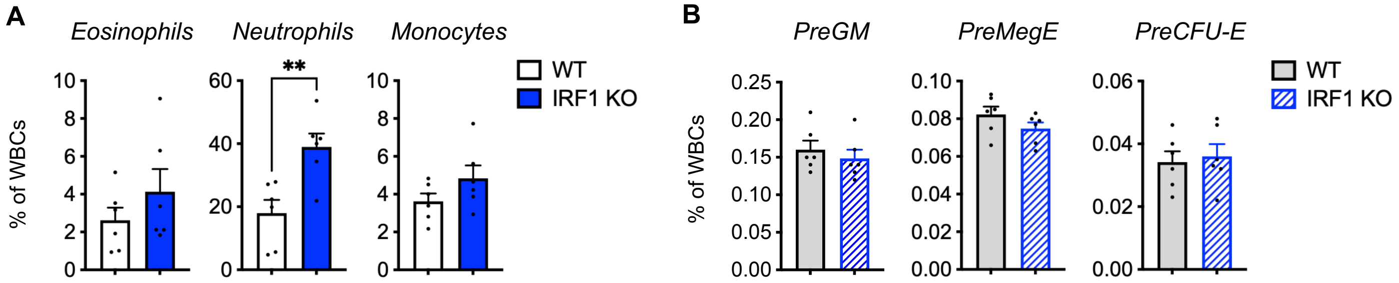

**Supplementary Figure 2.**

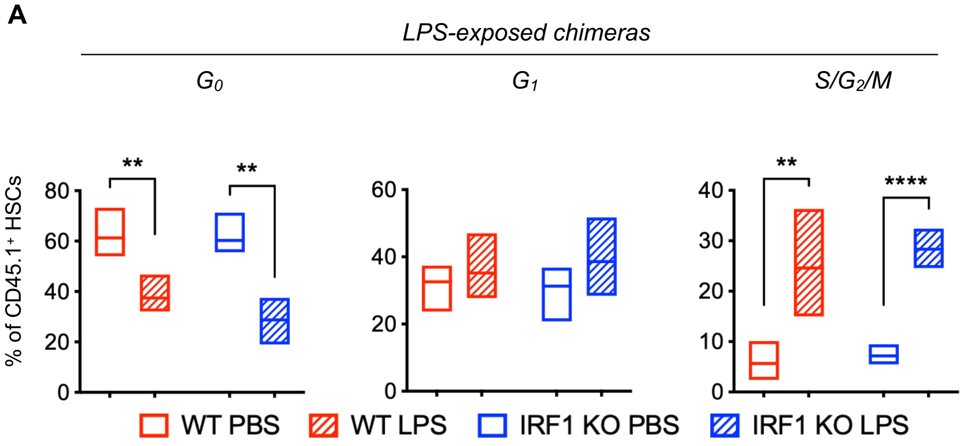

**Supplementary Figure 3.**

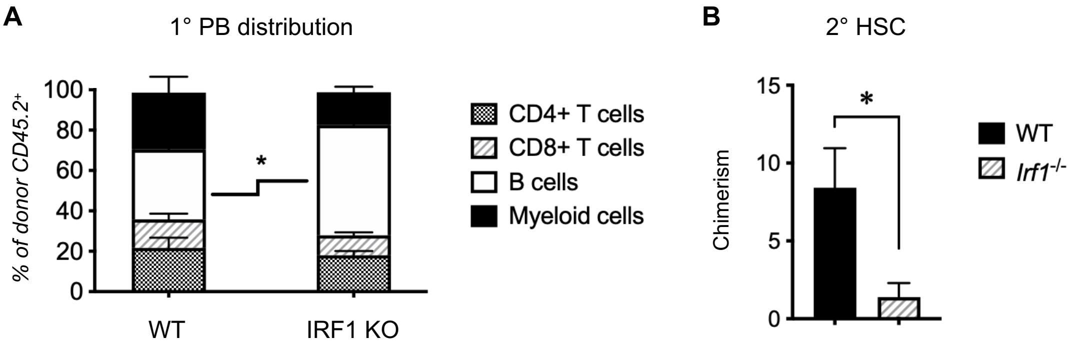

**Supplementary Figure 4.**

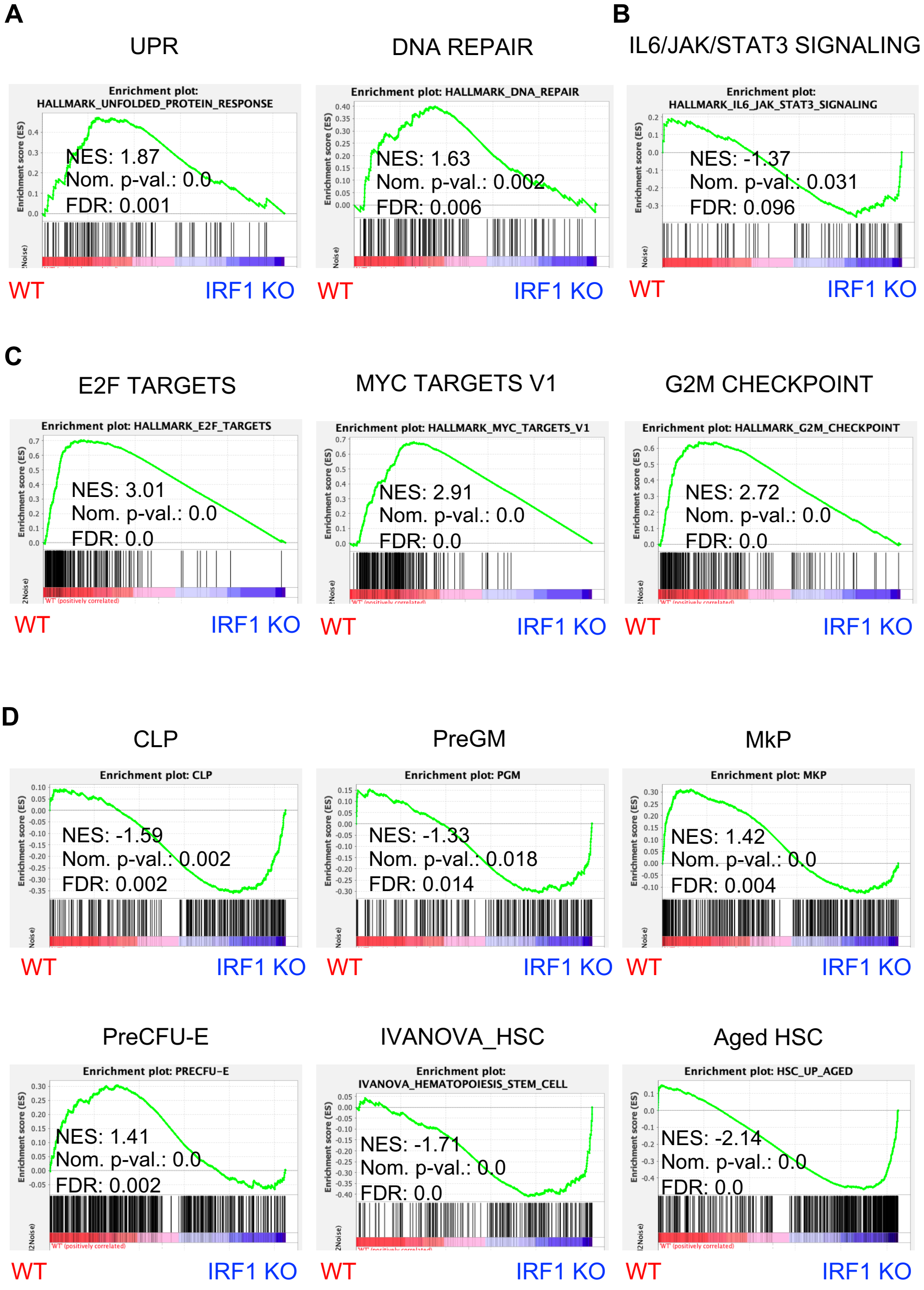

**Supplementary Figure 5.**

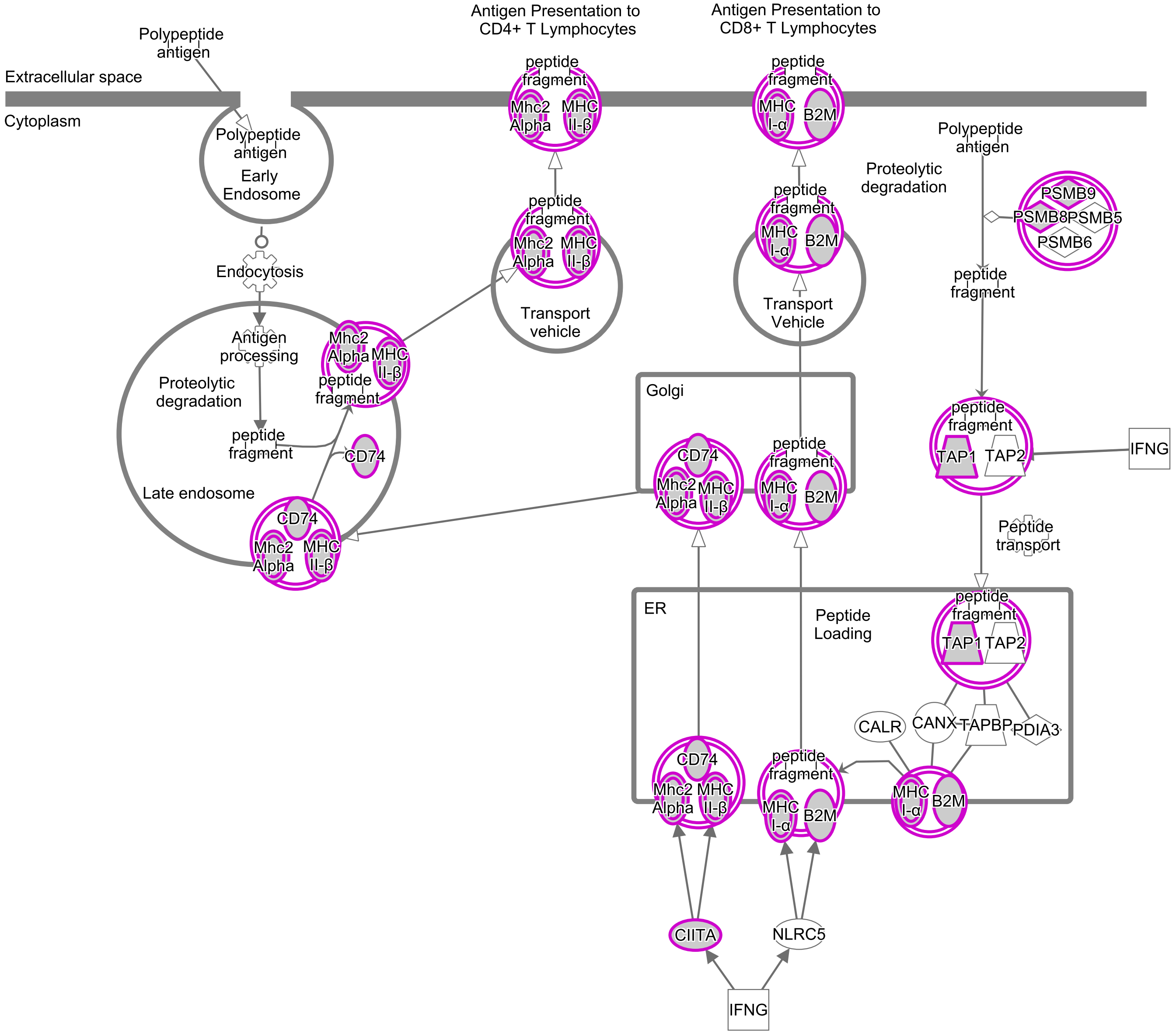

**Supplementary Figure 6.**

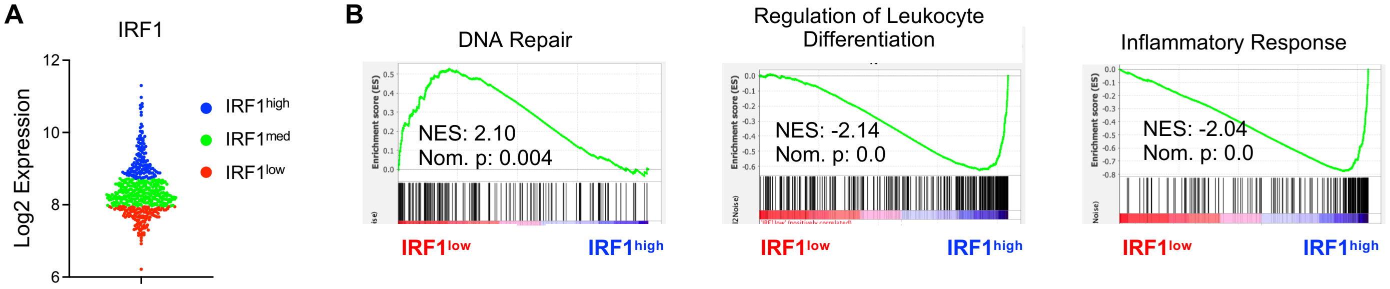

**Supplementary Figure 7.**

Supplementary Table 1 Downregulated genes in IRF1 KO HSCs

| GeneID | Base mean | log2(FC) | StdErr | Wald-Stats | P-value | P-adj |
|---|---|---|---|---|---|---|
| Apol7e | 417,7376 | 4,3988 | 0,1717 | 25,6223 | 0,0000 | 0,0000 |
| H2-Aa | 1 519,3001 | 3,9965 | 0,1518 | 26,3198 | 0,0000 | 0,0000 |
| H2-Eb1 | 725,9006 | 3,6804 | 0,1613 | 22,8155 | 0,0000 | 0,0000 |
| Zbp1 | 620,9153 | 3,5970 | 0,1551 | 23,1890 | 0,0000 | 0,0000 |
| Gm12185 | 877,2260 | 3,3470 | 0,1408 | 23,7662 | 0,0000 | 0,0000 |
| Tgtp1 | 2 283,7645 | 2,9696 | 0,1165 | 25,4954 | 0,0000 | 0,0000 |
| Gm4841 | 259,8222 | 2,7099 | 0,1706 | 15,8800 | 0,0000 | 0,0000 |
| AW112010 | 2 345,7773 | 2,6845 | 0,1089 | 24,6494 | 0,0000 | 0,0000 |
| Iigp1 | 2 783,7997 | 2,4268 | 0,1132 | 21,4358 | 0,0000 | 0,0000 |
| Gbp10 | 211,4595 | 2,4268 | 0,1820 | 13,3346 | 0,0000 | 0,0000 |
| Gm4951 | 923,5184 | 2,3398 | 0,1241 | 18,8537 | 0,0000 | 0,0000 |
| F830016B08Rik | 302,2336 | 2,2813 | 0,1738 | 13,1290 | 0,0000 | 0,0000 |
| Aspa | 235,1483 | 2,2681 | 0,1674 | 13,5524 | 0,0000 | 0,0000 |
| Casp1 | 1 181,0670 | 2,2502 | 0,1284 | 17,5257 | 0,0000 | 0,0000 |
| Casp12 | 1 107,0685 | 2,0632 | 0,1348 | 15,3111 | 0,0000 | 0,0000 |
| Il18bp | 577,5839 | 2,0503 | 0,1457 | 14,0699 | 0,0000 | 0,0000 |
| Gm12250 | 1 877,9331 | 1,8219 | 0,1820 | 10,0127 | 0,0000 | 0,0000 |
| Il12rb1 | 766,8363 | 1,7836 | 0,1576 | 11,3182 | 0,0000 | 0,0000 |
| Slamf8 | 106,6277 | 1,7803 | 0,1843 | 9,6573 | 0,0000 | 0,0000 |
| Art2b | 166,7998 | 1,5951 | 0,1805 | 8,8374 | 0,0000 | 0,0000 |
| Sfrp1 | 511,3875 | 1,5843 | 0,1489 | 10,6431 | 0,0000 | 0,0000 |
| H2-Ab1 | 3 574,9573 | 1,5442 | 0,1804 | 8,5592 | 0,0000 | 0,0000 |
| Apol9b | 78,9081 | 1,5082 | 0,1831 | 8,2392 | 0,0000 | 0,0000 |
| Aldh1a1 | 3 046,7934 | 1,4955 | 0,1018 | 14,6942 | 0,0000 | 0,0000 |
| Insl6 | 391,4946 | 1,4842 | 0,1504 | 9,8663 | 0,0000 | 0,0000 |
| Ifit1bl1 | 236,4974 | 1,4553 | 0,1820 | 7,9973 | 0,0000 | 0,0000 |
| Gbp4 | 1 165,4715 | 1,4529 | 0,1841 | 7,8938 | 0,0000 | 0,0000 |
| Phf11d | 250,0603 | 1,4205 | 0,1642 | 8,6504 | 0,0000 | 0,0000 |
| Itgb7 | 1 414,0180 | 1,4045 | 0,1428 | 9,8358 | 0,0000 | 0,0000 |
| Gbp8 | 1 067,1148 | 1,3537 | 0,1157 | 11,7020 | 0,0000 | 0,0000 |
| Il17rc | 169,2101 | 1,3126 | 0,1758 | 7,4678 | 0,0000 | 0,0000 |
| Rhbg | 90,0281 | 1,2995 | 0,1844 | 7,0472 | 0,0000 | 0,0000 |
| Ifit3 | 328,6710 | 1,2794 | 0,1689 | 7,5740 | 0,0000 | 0,0000 |
| Cysltr2 | 390,7980 | 1,2732 | 0,1497 | 8,5062 | 0,0000 | 0,0000 |
| Tapbpl | 3 055,2726 | 1,2272 | 0,0993 | 12,3641 | 0,0000 | 0,0000 |
| Insrr | 524,9477 | 1,2231 | 0,1639 | 7,4623 | 0,0000 | 0,0000 |
| Ifit3b | 109,6780 | 1,2022 | 0,1831 | 6,5655 | 0,0000 | 0,0000 |
| Gbp2 | 3 025,4706 | 1,1615 | 0,0936 | 12,4055 | 0,0000 | 0,0000 |
| Cd86 | 470,1546 | 1,1570 | 0,1381 | 8,3776 | 0,0000 | 0,0000 |
| Phactr1 | 629,5696 | 1,1305 | 0,1349 | 8,3811 | 0,0000 | 0,0000 |
| H2-Q6 | 355,9135 | 1,1269 | 0,1613 | 6,9878 | 0,0000 | 0,0000 |
| Oas3 | 1 836,2010 | 1,1117 | 0,1264 | 8,7926 | 0,0000 | 0,0000 |
| Stxbp4 | 1 861,5173 | 1,0926 | 0,1099 | 9,9406 | 0,0000 | 0,0000 |
| Ifi47 | 8 879,1261 | 1,0688 | 0,0942 | 11,3439 | 0,0000 | 0,0000 |
| Rhobtb1 | 1 118,4279 | 1,0421 | 0,1202 | 8,6713 | 0,0000 | 0,0000 |
| Dio2 | 311,1355 | 1,0382 | 0,1569 | 6,6187 | 0,0000 | 0,0000 |

| | | | | | | |
|---|---|---|---|---|---|---|
| Gm9199 | 254,3811 | 1,0121 | 0,1648 | 6,1429 | 0,0000 | 0,0000 |
| A930033H14Rik | 90,0775 | 1,0004 | 0,1842 | 5,4314 | 0,0000 | 0,0000 |
| Ciita | 374,5265 | 0,9988 | 0,1697 | 5,8870 | 0,0000 | 0,0000 |
| H2-T3 | 60,6938 | 0,9834 | 0,1789 | 5,4963 | 0,0000 | 0,0000 |
| Kalrn | 975,1803 | 0,9784 | 0,1398 | 6,9999 | 0,0000 | 0,0000 |
| BC064078 | 489,6560 | 0,9771 | 0,1407 | 6,9472 | 0,0000 | 0,0000 |
| Tcam1 | 785,0315 | 0,9661 | 0,1528 | 6,3216 | 0,0000 | 0,0000 |
| Cap2 | 169,1205 | 0,9583 | 0,1755 | 5,4621 | 0,0000 | 0,0000 |
| Gm1966 | 1 109,1325 | 0,9533 | 0,1345 | 7,0870 | 0,0000 | 0,0000 |
| Gm5431 | 92,2511 | 0,9495 | 0,1844 | 5,1494 | 0,0000 | 0,0000 |
| Lyrm7 | 245,8109 | 0,9457 | 0,1660 | 5,6983 | 0,0000 | 0,0000 |
| Alox5 | 369,4358 | 0,9384 | 0,1683 | 5,5751 | 0,0000 | 0,0000 |
| Tgtp2 | 6 024,6139 | 0,9298 | 0,1200 | 7,7500 | 0,0000 | 0,0000 |
| Psmb9 | 7 435,3359 | 0,9241 | 0,0834 | 11,0850 | 0,0000 | 0,0000 |
| Slco2a1 | 128,0752 | 0,9183 | 0,1827 | 5,0254 | 0,0000 | 0,0000 |
| Gbp6 | 1 714,7027 | 0,9177 | 0,1820 | 5,0430 | 0,0000 | 0,0000 |
| Batf2 | 693,4981 | 0,9138 | 0,1380 | 6,6220 | 0,0000 | 0,0000 |
| Tuba8 | 3 997,7985 | 0,9076 | 0,1159 | 7,8318 | 0,0000 | 0,0000 |
| Gm12216 | 553,0912 | 0,9012 | 0,1374 | 6,5574 | 0,0000 | 0,0000 |
| Adgrd1 | 1 442,2489 | 0,8972 | 0,1099 | 8,1670 | 0,0000 | 0,0000 |
| 9530082P21Rik | 246,7975 | 0,8937 | 0,1626 | 5,4951 | 0,0000 | 0,0000 |
| Psmb10 | 6 725,8782 | 0,8818 | 0,0974 | 9,0514 | 0,0000 | 0,0000 |
| Psme2b | 1 328,8789 | 0,8794 | 0,1161 | 7,5770 | 0,0000 | 0,0000 |
| B3galt1 | 87,0293 | 0,8769 | 0,1842 | 4,7599 | 0,0000 | 0,0001 |
| Gm43305 | 405,2747 | 0,8481 | 0,1676 | 5,0598 | 0,0000 | 0,0000 |
| Igtp | 5 940,1597 | 0,8309 | 0,1163 | 7,1418 | 0,0000 | 0,0000 |
| BC147527 | 535,8349 | 0,8303 | 0,1500 | 5,5371 | 0,0000 | 0,0000 |
| Naip6 | 86,0514 | 0,8299 | 0,1843 | 4,5023 | 0,0000 | 0,0003 |
| Psmb8 | 19 577,6351 | 0,8040 | 0,0933 | 8,6156 | 0,0000 | 0,0000 |
| Islr | 701,2857 | 0,7862 | 0,1458 | 5,3944 | 0,0000 | 0,0000 |
| Psme2 | 4 580,7820 | 0,7827 | 0,0946 | 8,2780 | 0,0000 | 0,0000 |
| Apol9a | 57,4986 | 0,7694 | 0,1816 | 4,2379 | 0,0000 | 0,0009 |
| Glrp1 | 128,3431 | 0,7689 | 0,1837 | 4,1853 | 0,0000 | 0,0010 |
| Ripply3 | 178,7155 | 0,7670 | 0,1754 | 4,3728 | 0,0000 | 0,0005 |
| Gp1bb | 1 654,7139 | 0,7661 | 0,1277 | 5,9998 | 0,0000 | 0,0000 |
| Stat1 | 12 776,8606 | 0,7611 | 0,1240 | 6,1355 | 0,0000 | 0,0000 |
| Sntb1 | 253,7972 | 0,7498 | 0,1566 | 4,7877 | 0,0000 | 0,0001 |
| Mx1 | 150,9120 | 0,7362 | 0,1803 | 4,0834 | 0,0000 | 0,0015 |
| Gbp3 | 2 040,1605 | 0,7299 | 0,1819 | 4,0123 | 0,0001 | 0,0019 |
| Adgrg7 | 232,6703 | 0,7287 | 0,1637 | 4,4503 | 0,0000 | 0,0004 |
| Erap1 | 7 164,9187 | 0,7286 | 0,0895 | 8,1368 | 0,0000 | 0,0000 |
| Ache | 1 690,4629 | 0,7166 | 0,1441 | 4,9733 | 0,0000 | 0,0000 |
| Gins1 | 735,8977 | 0,7125 | 0,1237 | 5,7612 | 0,0000 | 0,0000 |
| A530040E14Rik | 96,6006 | 0,7074 | 0,1834 | 3,8576 | 0,0001 | 0,0031 |
| Trim58 | 573,0748 | 0,7063 | 0,1637 | 4,3148 | 0,0000 | 0,0006 |
| E2f8 | 1 803,7994 | 0,7041 | 0,1264 | 5,5725 | 0,0000 | 0,0000 |
| Il7 | 220,8454 | 0,6937 | 0,1779 | 3,8985 | 0,0001 | 0,0027 |
| Gpc1 | 1 400,2035 | 0,6903 | 0,1056 | 6,5349 | 0,0000 | 0,0000 |

| | | | | | | |
|---|---|---|---|---|---|---|
| Scly | 1 877,7366 | 0,6854 | 0,1049 | 6,5365 | 0,0000 | 0,0000 |
| Phf11b | 539,6903 | 0,6828 | 0,1565 | 4,3630 | 0,0000 | 0,0005 |
| Raver2 | 1 029,6555 | 0,6821 | 0,1121 | 6,0842 | 0,0000 | 0,0000 |
| Pf4 | 5 418,8746 | 0,6781 | 0,1765 | 3,8412 | 0,0001 | 0,0032 |
| Trafd1 | 4 256,8036 | 0,6763 | 0,0868 | 7,7944 | 0,0000 | 0,0000 |
| Treml1 | 1 508,2286 | 0,6736 | 0,1320 | 5,1029 | 0,0000 | 0,0000 |
| Ifi27 | 2 941,9431 | 0,6723 | 0,1404 | 4,7875 | 0,0000 | 0,0001 |
| Zdhhc14 | 432,9322 | 0,6630 | 0,1540 | 4,3061 | 0,0000 | 0,0007 |
| Parm1 | 229,1493 | 0,6612 | 0,1720 | 3,8449 | 0,0001 | 0,0032 |
| Nlrp1a | 265,0472 | 0,6589 | 0,1599 | 4,1211 | 0,0000 | 0,0013 |
| Knl1 | 1 414,1730 | 0,6535 | 0,1302 | 5,0186 | 0,0000 | 0,0000 |
| Isg20 | 1 976,2844 | 0,6527 | 0,1077 | 6,0579 | 0,0000 | 0,0000 |
| Melk | 1 560,7813 | 0,6512 | 0,1403 | 4,6409 | 0,0000 | 0,0002 |
| AI504432 | 125,1559 | 0,6463 | 0,1789 | 3,6114 | 0,0003 | 0,0066 |
| Ly75 | 1 677,9091 | 0,6397 | 0,1020 | 6,2685 | 0,0000 | 0,0000 |
| Top2a | 9 528,4307 | 0,6382 | 0,1310 | 4,8706 | 0,0000 | 0,0001 |
| Ifit2 | 3 011,0220 | 0,6379 | 0,0944 | 6,7586 | 0,0000 | 0,0000 |
| H2-DMa | 7 930,6021 | 0,6359 | 0,1056 | 6,0231 | 0,0000 | 0,0000 |
| Hdac11 | 409,1003 | 0,6313 | 0,1423 | 4,4350 | 0,0000 | 0,0004 |
| H2-Q5 | 2 400,8998 | 0,6273 | 0,1374 | 4,5653 | 0,0000 | 0,0002 |
| Kcna3 | 2 581,8869 | 0,6269 | 0,1203 | 5,2103 | 0,0000 | 0,0000 |
| Nuak2 | 1 352,1206 | 0,6223 | 0,1079 | 5,7696 | 0,0000 | 0,0000 |
| Ddx60 | 1 281,4083 | 0,6193 | 0,1120 | 5,5276 | 0,0000 | 0,0000 |
| Fbn1 | 158,5692 | 0,6150 | 0,1836 | 3,3496 | 0,0008 | 0,0138 |
| Pbk | 4 209,0587 | 0,6135 | 0,1279 | 4,7951 | 0,0000 | 0,0001 |
| Hrh2 | 350,5639 | 0,6108 | 0,1637 | 3,7321 | 0,0002 | 0,0046 |
| Crybg1 | 537,9201 | 0,6101 | 0,1468 | 4,1548 | 0,0000 | 0,0011 |
| Gp1ba | 1 434,2978 | 0,6070 | 0,1297 | 4,6817 | 0,0000 | 0,0001 |
| Sphk1 | 109,0338 | 0,6046 | 0,1761 | 3,4344 | 0,0006 | 0,0109 |
| Plek | 8 857,7158 | 0,5995 | 0,1801 | 3,3294 | 0,0009 | 0,0145 |
| Cd74 | 3 314,6070 | 0,5975 | 0,1433 | 4,1702 | 0,0000 | 0,0011 |
| Cldn10 | 105,0351 | 0,5951 | 0,1838 | 3,2383 | 0,0012 | 0,0186 |
| Slc22a3 | 6 034,9294 | 0,5951 | 0,1128 | 5,2752 | 0,0000 | 0,0000 |
| B2m | 82 577,8316 | 0,5947 | 0,0767 | 7,7555 | 0,0000 | 0,0000 |
| Asb13 | 2 397,9108 | 0,5903 | 0,1110 | 5,3169 | 0,0000 | 0,0000 |
| Tap1 | 15 163,5860 | 0,5871 | 0,1005 | 5,8392 | 0,0000 | 0,0000 |
| Rap1b | 29 851,8350 | 0,5868 | 0,1087 | 5,3986 | 0,0000 | 0,0000 |
| F2r | 14 332,4167 | 0,5864 | 0,1222 | 4,7973 | 0,0000 | 0,0001 |
| Irgm2 | 5 003,9050 | 0,5848 | 0,1042 | 5,6146 | 0,0000 | 0,0000 |
| Mdm1 | 1 416,9382 | 0,5844 | 0,1466 | 3,9865 | 0,0001 | 0,0020 |

Supplementary Table 2 Upregulated genes in IRF1 KO HSCs

| GeneID | Base mean | log2(FC) | StdErr | Wald-Stats | P-value | P-adj |
|---|---|---|---|---|---|---|
| Ehd2 | 328,2852 | -1,4571 | 0,1557 | -9,3613 | 0,0000 | 0,0000 |
| Smoc1 | 225,6872 | -1,2158 | 0,1770 | -6,8697 | 0,0000 | 0,0000 |
| Il10ra | 412,6328 | -1,1297 | 0,1639 | -6,8911 | 0,0000 | 0,0000 |
| Pim2 | 9 974,9428 | -1,0726 | 0,1270 | -8,4443 | 0,0000 | 0,0000 |
| Lamb2 | 1 288,1787 | -1,0310 | 0,1466 | -7,0324 | 0,0000 | 0,0000 |
| Rab34 | 259,9217 | -1,0212 | 0,1619 | -6,3064 | 0,0000 | 0,0000 |
| Mapk13 | 164,4534 | -1,0165 | 0,1715 | -5,9273 | 0,0000 | 0,0000 |
| Sparc | 210,2071 | -1,0024 | 0,1794 | -5,5866 | 0,0000 | 0,0000 |
| C4b | 257,1744 | -0,9944 | 0,1824 | -5,4529 | 0,0000 | 0,0000 |
| Socs3 | 1 163,2932 | -0,9842 | 0,1333 | -7,3853 | 0,0000 | 0,0000 |
| Phldb2 | 727,5672 | -0,9795 | 0,1320 | -7,4211 | 0,0000 | 0,0000 |
| Col4a6 | 78,3953 | -0,9704 | 0,1843 | -5,2646 | 0,0000 | 0,0000 |
| Mmp2 | 1 303,3885 | -0,9673 | 0,1551 | -6,2350 | 0,0000 | 0,0000 |
| Cyp27a1 | 674,1308 | -0,9591 | 0,1513 | -6,3400 | 0,0000 | 0,0000 |
| Flrt3 | 176,3989 | -0,9555 | 0,1778 | -5,3752 | 0,0000 | 0,0000 |
| 1700003E16Rik | 358,0102 | -0,9513 | 0,1484 | -6,4105 | 0,0000 | 0,0000 |
| Grrp1 | 120,5051 | -0,9358 | 0,1840 | -5,0860 | 0,0000 | 0,0000 |
| Egr1 | 4 401,3766 | -0,9354 | 0,0975 | -9,5889 | 0,0000 | 0,0000 |
| Cd53 | 785,8289 | -0,9116 | 0,1512 | -6,0274 | 0,0000 | 0,0000 |
| Rap1gap | 737,6686 | -0,8946 | 0,1530 | -5,8460 | 0,0000 | 0,0000 |
| Arhgef25 | 806,7774 | -0,8945 | 0,1587 | -5,6377 | 0,0000 | 0,0000 |
| Scube3 | 252,1784 | -0,8856 | 0,1717 | -5,1593 | 0,0000 | 0,0000 |
| Cd209f | 165,4274 | -0,8823 | 0,1835 | -4,8086 | 0,0000 | 0,0001 |
| Slc16a11 | 160,6015 | -0,8816 | 0,1823 | -4,8351 | 0,0000 | 0,0001 |
| Luzp2 | 102,5188 | -0,8758 | 0,1834 | -4,7755 | 0,0000 | 0,0001 |
| Itgb5 | 821,4674 | -0,8641 | 0,1322 | -6,5356 | 0,0000 | 0,0000 |
| Arid5a | 3 212,2126 | -0,8633 | 0,1069 | -8,0758 | 0,0000 | 0,0000 |
| Epcam | 314,7979 | -0,8577 | 0,1662 | -5,1601 | 0,0000 | 0,0000 |
| Meg3 | 375,1182 | -0,8552 | 0,1721 | -4,9690 | 0,0000 | 0,0000 |
| Bmp4 | 166,6810 | -0,8446 | 0,1784 | -4,7356 | 0,0000 | 0,0001 |
| Ramp2 | 255,6162 | -0,8330 | 0,1716 | -4,8543 | 0,0000 | 0,0001 |
| Rorc | 548,4850 | -0,8311 | 0,1512 | -5,4970 | 0,0000 | 0,0000 |
| Snx31 | 453,5424 | -0,8251 | 0,1669 | -4,9426 | 0,0000 | 0,0000 |
| Tspan8 | 324,8225 | -0,8227 | 0,1675 | -4,9117 | 0,0000 | 0,0001 |
| Acox2 | 208,1272 | -0,8216 | 0,1736 | -4,7332 | 0,0000 | 0,0001 |
| Tnfrsf25 | 230,3779 | -0,8210 | 0,1661 | -4,9427 | 0,0000 | 0,0000 |
| Gm5833 | 204,5609 | -0,8172 | 0,1779 | -4,5921 | 0,0000 | 0,0002 |
| Tmem150b | 78,3348 | -0,8126 | 0,1843 | -4,4088 | 0,0000 | 0,0004 |
| Car11 | 232,8623 | -0,8116 | 0,1764 | -4,6000 | 0,0000 | 0,0002 |
| Pdk4 | 311,0985 | -0,8081 | 0,1628 | -4,9646 | 0,0000 | 0,0000 |
| Cebpd | 312,8182 | -0,8077 | 0,1535 | -5,2605 | 0,0000 | 0,0000 |
| Efnb2 | 401,3571 | -0,8072 | 0,1548 | -5,2129 | 0,0000 | 0,0000 |
| Adamtsl4 | 268,1057 | -0,8060 | 0,1799 | -4,4802 | 0,0000 | 0,0003 |
| Fam181b | 895,1355 | -0,7954 | 0,1391 | -5,7174 | 0,0000 | 0,0000 |
| Arap3 | 932,5481 | -0,7929 | 0,1197 | -6,6245 | 0,0000 | 0,0000 |
| Vldlr | 3 087,2049 | -0,7913 | 0,1295 | -6,1081 | 0,0000 | 0,0000 |

| | | | | | | |
|---|---|---|---|---|---|---|
| Pld4 | 343,3543 | -0,7858 | 0,1792 | -4,3859 | 0,0000 | 0,0005 |
| Tgfbi | 1 891,5851 | -0,7839 | 0,1122 | -6,9867 | 0,0000 | 0,0000 |
| Susd4 | 173,9416 | -0,7836 | 0,1721 | -4,5522 | 0,0000 | 0,0002 |
| Fgd5 | 5 213,1557 | -0,7804 | 0,0877 | -8,8963 | 0,0000 | 0,0000 |
| Ldb2 | 207,0271 | -0,7780 | 0,1734 | -4,4862 | 0,0000 | 0,0003 |
| Dll4 | 217,9321 | -0,7732 | 0,1683 | -4,5942 | 0,0000 | 0,0002 |
| Rin2 | 1 221,7270 | -0,7695 | 0,1355 | -5,6796 | 0,0000 | 0,0000 |
| Bcam | 1 331,3760 | -0,7619 | 0,1203 | -6,3359 | 0,0000 | 0,0000 |
| Lsr | 1 241,0691 | -0,7575 | 0,1212 | -6,2499 | 0,0000 | 0,0000 |
| Selp | 4 628,1621 | -0,7508 | 0,1194 | -6,2887 | 0,0000 | 0,0000 |
| Fkbp10 | 551,2896 | -0,7498 | 0,1497 | -5,0099 | 0,0000 | 0,0000 |
| Gfi1 | 1 033,4080 | -0,7497 | 0,1467 | -5,1114 | 0,0000 | 0,0000 |
| Rftn2 | 160,3391 | -0,7473 | 0,1808 | -4,1344 | 0,0000 | 0,0012 |
| Baiap3 | 82,7507 | -0,7419 | 0,1774 | -4,1816 | 0,0000 | 0,0011 |
| Il18rap | 1 214,5949 | -0,7381 | 0,1418 | -5,2054 | 0,0000 | 0,0000 |
| Hk3 | 2 996,7362 | -0,7369 | 0,1139 | -6,4669 | 0,0000 | 0,0000 |
| Ptges | 1 041,2381 | -0,7343 | 0,1304 | -5,6301 | 0,0000 | 0,0000 |
| Dpp4 | 4 242,5344 | -0,7330 | 0,1059 | -6,9206 | 0,0000 | 0,0000 |
| Pbld1 | 474,7476 | -0,7325 | 0,1662 | -4,4080 | 0,0000 | 0,0004 |
| Nectin2 | 336,2803 | -0,7238 | 0,1689 | -4,2849 | 0,0000 | 0,0007 |
| Bcl3 | 2 383,3791 | -0,7222 | 0,1157 | -6,2405 | 0,0000 | 0,0000 |
| P2rx6 | 300,9663 | -0,7178 | 0,1637 | -4,3848 | 0,0000 | 0,0005 |
| Rab44 | 308,5739 | -0,7157 | 0,1580 | -4,5312 | 0,0000 | 0,0003 |
| Prom2 | 313,9157 | -0,7073 | 0,1818 | -3,8913 | 0,0001 | 0,0028 |
| Rab36 | 495,7760 | -0,7051 | 0,1382 | -5,1013 | 0,0000 | 0,0000 |
| Tubg2 | 67,1766 | -0,6947 | 0,1821 | -3,8159 | 0,0001 | 0,0035 |
| Clec4e | 660,2620 | -0,6940 | 0,1429 | -4,8573 | 0,0000 | 0,0001 |
| Efna1 | 1 559,7049 | -0,6907 | 0,1303 | -5,2999 | 0,0000 | 0,0000 |
| 1810073O08Rik | 2 685,4954 | -0,6905 | 0,1249 | -5,5263 | 0,0000 | 0,0000 |
| Rnf208 | 609,6488 | -0,6900 | 0,1493 | -4,6215 | 0,0000 | 0,0002 |
| Krt80 | 447,1023 | -0,6895 | 0,1516 | -4,5490 | 0,0000 | 0,0003 |
| Agtr1a | 121,7336 | -0,6884 | 0,1803 | -3,8185 | 0,0001 | 0,0035 |
| Laptm4b | 2 608,4482 | -0,6868 | 0,1042 | -6,5919 | 0,0000 | 0,0000 |
| Veph1 | 169,7086 | -0,6849 | 0,1741 | -3,9331 | 0,0001 | 0,0024 |
| Efemp2 | 170,7798 | -0,6838 | 0,1829 | -3,7388 | 0,0002 | 0,0045 |
| Csf2rb | 2 584,7768 | -0,6823 | 0,1332 | -5,1232 | 0,0000 | 0,0000 |
| Sult1a1 | 3 818,8282 | -0,6802 | 0,1272 | -5,3471 | 0,0000 | 0,0000 |
| Pld2 | 1 205,0355 | -0,6781 | 0,1305 | -5,1957 | 0,0000 | 0,0000 |
| Klc3 | 307,2066 | -0,6763 | 0,1602 | -4,2204 | 0,0000 | 0,0009 |
| Nfil3 | 334,6085 | -0,6756 | 0,1459 | -4,6309 | 0,0000 | 0,0002 |
| Fosl2 | 260,3871 | -0,6738 | 0,1616 | -4,1693 | 0,0000 | 0,0011 |
| Lama5 | 286,8845 | -0,6730 | 0,1821 | -3,6962 | 0,0002 | 0,0051 |
| Nox4 | 91,7056 | -0,6696 | 0,1839 | -3,6417 | 0,0003 | 0,0060 |
| Vwa7 | 129,2156 | -0,6659 | 0,1841 | -3,6168 | 0,0003 | 0,0065 |
| Kifc2 | 350,9118 | -0,6644 | 0,1523 | -4,3615 | 0,0000 | 0,0005 |
| Tdrd9 | 168,1670 | -0,6632 | 0,1735 | -3,8218 | 0,0001 | 0,0034 |
| St8sia4 | 2 916,8958 | -0,6631 | 0,1000 | -6,6326 | 0,0000 | 0,0000 |
| Enho | 481,9614 | -0,6631 | 0,1521 | -4,3587 | 0,0000 | 0,0005 |

| | | | | | | |
|---|---|---|---|---|---|---|
| Dcaf12l2 | 84,7988 | -0,6630 | 0,1836 | -3,6122 | 0,0003 | 0,0066 |
| Spint1 | 119,6477 | -0,6629 | 0,1838 | -3,6070 | 0,0003 | 0,0067 |
| Fgfr3 | 1 237,9654 | -0,6566 | 0,1300 | -5,0518 | 0,0000 | 0,0000 |
| Tmem44 | 361,7029 | -0,6547 | 0,1701 | -3,8478 | 0,0001 | 0,0032 |
| Wfdc2 | 586,5905 | -0,6534 | 0,1445 | -4,5221 | 0,0000 | 0,0003 |
| Lrtm2 | 265,2716 | -0,6533 | 0,1652 | -3,9557 | 0,0001 | 0,0022 |
| Ifitm1 | 26 195,8619 | -0,6524 | 0,1230 | -5,3035 | 0,0000 | 0,0000 |
| Spats2 | 201,8436 | -0,6493 | 0,1780 | -3,6474 | 0,0003 | 0,0059 |
| Rnf39 | 92,0649 | -0,6458 | 0,1818 | -3,5520 | 0,0004 | 0,0079 |
| Sema4f | 158,7213 | -0,6439 | 0,1844 | -3,4922 | 0,0005 | 0,0092 |
| Plekhb1 | 230,3010 | -0,6414 | 0,1621 | -3,9573 | 0,0001 | 0,0022 |
| Dnm3 | 96,5122 | -0,6401 | 0,1844 | -3,4716 | 0,0005 | 0,0098 |
| Cd14 | 98,5125 | -0,6398 | 0,1835 | -3,4856 | 0,0005 | 0,0094 |
| Smarca1 | 195,1537 | -0,6390 | 0,1778 | -3,5932 | 0,0003 | 0,0070 |
| Wfdc17 | 193,4139 | -0,6390 | 0,1830 | -3,4919 | 0,0005 | 0,0092 |
| Ctsh | 1 152,0890 | -0,6385 | 0,1068 | -5,9764 | 0,0000 | 0,0000 |
| Sec31b | 468,4440 | -0,6378 | 0,1358 | -4,6972 | 0,0000 | 0,0001 |
| Kif19a | 96,6640 | -0,6372 | 0,1844 | -3,4559 | 0,0005 | 0,0103 |
| Il1r1 | 978,7910 | -0,6350 | 0,1370 | -4,6348 | 0,0000 | 0,0002 |
| Kcnd1 | 529,1039 | -0,6340 | 0,1529 | -4,1455 | 0,0000 | 0,0012 |
| Slc26a11 | 1 407,5738 | -0,6316 | 0,1244 | -5,0777 | 0,0000 | 0,0000 |
| Fgf11 | 682,8206 | -0,6306 | 0,1348 | -4,6770 | 0,0000 | 0,0001 |
| Rcan2 | 123,1996 | -0,6292 | 0,1802 | -3,4910 | 0,0005 | 0,0092 |
| Csf2rb2 | 599,8189 | -0,6227 | 0,1571 | -3,9641 | 0,0001 | 0,0022 |
| Tifab | 104,0146 | -0,6210 | 0,1835 | -3,3839 | 0,0007 | 0,0127 |
| Sstr2 | 125,9527 | -0,6204 | 0,1835 | -3,3812 | 0,0007 | 0,0127 |
| Eps8 | 3 273,7375 | -0,6197 | 0,1033 | -5,9979 | 0,0000 | 0,0000 |
| Ppp1r3b | 2 426,3155 | -0,6193 | 0,1122 | -5,5205 | 0,0000 | 0,0000 |
| Plekhg6 | 524,9541 | -0,6188 | 0,1563 | -3,9582 | 0,0001 | 0,0022 |
| Slfn2 | 4 585,6400 | -0,6185 | 0,1144 | -5,4085 | 0,0000 | 0,0000 |
| Ppargc1a | 314,0484 | -0,6175 | 0,1667 | -3,7044 | 0,0002 | 0,0049 |
| Hid1 | 5 802,5304 | -0,6174 | 0,0999 | -6,1791 | 0,0000 | 0,0000 |
| Mmp14 | 1 942,7532 | -0,6161 | 0,1025 | -6,0110 | 0,0000 | 0,0000 |
| Cacna2d4 | 279,4213 | -0,6154 | 0,1628 | -3,7791 | 0,0002 | 0,0039 |
| Igdcc4 | 911,4583 | -0,6150 | 0,1444 | -4,2578 | 0,0000 | 0,0008 |
| Syne4 | 180,9783 | -0,6128 | 0,1750 | -3,5015 | 0,0005 | 0,0090 |
| Slc4a3 | 253,2828 | -0,6113 | 0,1705 | -3,5861 | 0,0003 | 0,0071 |
| Epas1 | 94,3195 | -0,6111 | 0,1839 | -3,3234 | 0,0009 | 0,0147 |
| Nhsl2 | 570,0408 | -0,6108 | 0,1531 | -3,9880 | 0,0001 | 0,0020 |
| Arhgef10l | 791,9478 | -0,6105 | 0,1459 | -4,1840 | 0,0000 | 0,0010 |
| Trim62 | 1 663,2585 | -0,6075 | 0,1269 | -4,7868 | 0,0000 | 0,0001 |
| Acot1 | 696,7001 | -0,6069 | 0,1264 | -4,8020 | 0,0000 | 0,0001 |
| Il18r1 | 197,8274 | -0,6056 | 0,1803 | -3,3594 | 0,0008 | 0,0135 |
| S1pr1 | 3 100,5409 | -0,6025 | 0,1093 | -5,5122 | 0,0000 | 0,0000 |
| Rgl1 | 1 983,3969 | -0,6020 | 0,1080 | -5,5759 | 0,0000 | 0,0000 |
| Lrrn3 | 214,4210 | -0,6009 | 0,1771 | -3,3927 | 0,0007 | 0,0124 |
| Trim2 | 170,3840 | -0,6003 | 0,1740 | -3,4503 | 0,0006 | 0,0104 |
| Fam43a | 5 542,5252 | -0,5998 | 0,0884 | -6,7879 | 0,0000 | 0,0000 |

| | | | | | | |
|---|---|---|---|---|---|---|
| Cyp39a1 | 309,4980 | -0,5998 | 0,1515 | -3,9587 | 0,0001 | 0,0022 |
| Rasd1 | 121,7476 | -0,5994 | 0,1826 | -3,2835 | 0,0010 | 0,0163 |
| Sema6c | 589,4367 | -0,5991 | 0,1417 | -4,2268 | 0,0000 | 0,0009 |
| Wdr86 | 182,6591 | -0,5988 | 0,1841 | -3,2523 | 0,0011 | 0,0179 |
| Man1c1 | 2 341,9717 | -0,5977 | 0,1176 | -5,0846 | 0,0000 | 0,0000 |
| Trf | 5 432,3716 | -0,5956 | 0,0882 | -6,7522 | 0,0000 | 0,0000 |
| Kiss1r | 106,3283 | -0,5954 | 0,1837 | -3,2406 | 0,0012 | 0,0185 |
| Zfp185 | 140,4565 | -0,5954 | 0,1775 | -3,3542 | 0,0008 | 0,0137 |
| Tecpr1 | 3 869,3012 | -0,5946 | 0,1158 | -5,1327 | 0,0000 | 0,0000 |
| Msrb3 | 669,6692 | -0,5942 | 0,1691 | -3,5137 | 0,0004 | 0,0087 |
| Dlk1 | 303,3467 | -0,5917 | 0,1674 | -3,5350 | 0,0004 | 0,0082 |
| Prkar1b | 188,5284 | -0,5908 | 0,1844 | -3,2040 | 0,0014 | 0,0205 |
| Fcer1g | 1 785,4975 | -0,5902 | 0,1094 | -5,3942 | 0,0000 | 0,0000 |
| Tro | 793,6075 | -0,5892 | 0,1382 | -4,2619 | 0,0000 | 0,0008 |
| Cyp11a1 | 294,6479 | -0,5891 | 0,1578 | -3,7326 | 0,0002 | 0,0046 |
| Dsc2 | 243,8419 | -0,5879 | 0,1760 | -3,3396 | 0,0008 | 0,0142 |
| Pnck | 1 485,6861 | -0,5854 | 0,1085 | -5,3942 | 0,0000 | 0,0000 |
| Timp2 | 8 061,1999 | -0,5851 | 0,0832 | -7,0289 | 0,0000 | 0,0000 |
| Echdc2 | 222,4145 | -0,5842 | 0,1664 | -3,5102 | 0,0004 | 0,0088 |
| Mamdc2 | 3 936,6239 | -0,5838 | 0,0987 | -5,9124 | 0,0000 | 0,0000 |
| I730030J21Rik | 1 094,5387 | -0,5838 | 0,1214 | -4,8081 | 0,0000 | 0,0001 |
| Myo7a | 2 354,0769 | -0,5836 | 0,1128 | -5,1729 | 0,0000 | 0,0000 |
| Tiam2 | 423,5985 | -0,5836 | 0,1483 | -3,9343 | 0,0001 | 0,0024 |
| Zfr2 | 919,4336 | -0,5831 | 0,1296 | -4,5009 | 0,0000 | 0,0003 |
| Mcoln3 | 168,6191 | -0,5829 | 0,1799 | -3,2402 | 0,0012 | 0,0185 |
| Card6 | 217,1003 | -0,5824 | 0,1730 | -3,3671 | 0,0008 | 0,0132 |
| Lgals1 | 795,9438 | -0,5804 | 0,1482 | -3,9161 | 0,0001 | 0,0025 |

Supplementary Table 3 – IPA of downregulated genes in IRF1 KO HSCs

| Upstream Regulator | Molecule Type | p-value of overlap | Target Molecules in Dataset |
|---|---|---|---|
| STAT1 | transcription regulator | 1,43×10^-31 | B2M,BATF2,CASP1,CD86,CIITA,CYSLTR2,DDX60,GBP2,GBP4,GBP6,HLA-DQA1,HLA-DRB5,Ifi47,IFIT1,IFIT2,IFIT3,Igtp,Iigp1,IL18BP,Mx1,OAS3,PF4,PHACTR1,PSMB10,PSMB8,PSMB9,PSME2,SLAMF8,STAT1,TAP1,TAPBPL,Tgtp1/Tgtp2,TRAFD1,ZBP1 |
| IFNG | cytokine | 3,60×10^-30 | B2M,BATF2,CASP1,Casp12,CD74,CD86,CIITA,CYSLTR2,DDX60,ERAP1,F2R,GBP2,GBP4,GBP6,Gbp8,HLA-A,HLA-DMA,HLA-DQA1,HLA-DQB1,HLA-DRB5,Ifi47,IFIT1,IFIT2,IFIT3,Igtp,Iigp1,IL12RB1,IL18BP,IL7,INSRR,IRGM,ISG20,KNL1,LY75,Mx1,OAS3,PBK,PF4,PHACTR1,PHF11,PLEK,PSMB10,PSMB8,PSMB9,PSME2,SCLY,SFRP1,STAT1,TAP1,TAPBPL,Tgtp1/Tgtp2,TRAFD1 |
| lipopolysaccharide | chemical drug | 7,68×10^-29 | 9930111J21Rik1/Gm12185,ACHE,ALDH1A1,ALOX5,Apol7a (includes others),B2M,BC147527,CASP1,Casp12,CD74,CD86,CIITA,DDX60,DIO2,F2R,FBN1,GBP2,GBP4,GBP6,Gbp8,Gm12216,H2-Q5,HDAC11,HLA-A,HLA-DMA,HLA-DQA1,HLA-DQB1,HLA-DRB5,Ifi27,Ifi47,IFIT1,IFIT2,IFIT3,Igtp,Iigp1,IL12RB1,IL18BP,IL7,IRGM,ISG20,ITGB7,KCNA3,LY75,Mx1,OAS3,PBK,PF4,PHF11,PLEK,PSMB10,PSMB8,PSMB9,PSME2,SLAMF8,SLCO2A1,SPHK1,STAT1,TAP1,Tgtp1/Tgtp2,TOP2A,TRAFD1,ZBP1 |
| poly rI:rC-RNA | biologic drug | 1,80×10^-25 | Apol7a (includes others),Apol9a/Apol9b,Art2b,ASB13,B2M,Casp12,CD74,CD86,CIITA,DDX60,DIO2,GBP2,GBP4,GBP6,Gm4951,HLA-A,IFIT1,IFIT2,IFIT3,Igtp,Iigp1,IRGM,ISG20,Mx1,Naip1 (includes others),OAS3,PHF11,PSMB8,PSMB9,PSME2,STAT1,TAP1,TOP2A,TRAFD1,ZBP1 |
| Irgm1 | other | 2,16×10^-25 | Apol9a/Apol9b,B2M,CRYBG1,DDX60,GBP2,GBP4,GBP6,Gbp8,GINS1,Ifi27,IFIT2,IFIT3,Iigp1,IRGM,ISG20,Mx1,OAS3,STAT1,ZBP1 |
| TRIM24 | transcription regulator | 3,97×10^-24 | Apol9a/Apol9b,DDX60,FBN1,GBP2,GBP4,Ifi47,IFIT2,IFIT3,Igtp,Iigp1,IRGM,PHF11,PSMB10,PSMB8,PSMB9,STAT1,TAP1,Tgtp1/Tgtp2,TRAFD1 |
| IRF1 | transcription regulator | 1,03×10^-21 | B2M,CASP1,CIITA,ERAP1,GBP2,GBP4,Ifi47,IFIT1,IFIT2,IFIT3,IL12RB1,IL18BP,IL7,OAS3,PF4,PSMB10,PSMB8,PSMB9,PSME2,STAT1,TAP1 |
| Interferon alpha | group | 4,02×10^-21 | B2M,BATF2,CASP1,CD86,CIITA,F2R,GBP2,GBP6,HLA-A,Ifi27,Ifi47,IFIT1,IFIT2,IFIT3,Iigp1,IL12RB1,IL7,ISG20,Mx1,OAS3,PBK,PHF11,PSMB10,PSMB8,PSMB9,STAT1,TAP1,Tgtp1/Tgtp2,ZBP1 |
| USP8 | peptidase | 7,49×10^-21 | B2M,CASP1,ERAP1,HLA-A,IFIT1,IFIT2,IFIT3,IRGM,OAS3,PSMB10,PSMB8,PSMB9,STAT1,TAP1,TAPBPL |
| G protein alpha i | group | 1,84×10^-20 | F2R,GBP2,GBP4,GBP6,Gbp8,Gm4951,HLA-A,IFIT3,Igtp,Iigp1,IRGM,PSMB8,PSMB9,SPHK1,TAP1 |
| IRF7 | transcription regulator | 2,27×10^-20 | BC147527,GBP4,Ifi47,IFIT1,IFIT2,IFIT3,Igtp,IRGM,ISG20,Mx1,OAS3,PHF11,PSMB10,PSMB8,PSMB9,PSME2,STAT1,TAP1,ZBP1 |
| IFNB1 | cytokine | 1,11×10^-19 | Apol7a (includes others),CASP1,CD86,F2R,FBN1,GBP2,GBP4,GBP6,Gbp8,HLA-A,Ifi47,IFIT1,IFIT2,IFIT3,Igtp,IRGM,ISG20,Mx1,STAT1,Tgtp1/Tgtp2,ZBP1,ZDHHC14 |
| IL10RA | transmembrane receptor | 2,77×10^-19 | ALOX5,Art2b,BATF2,FBN1,GBP2,GBP6,Gbp8,HLA-A,Ifi47,Iigp1,IL12RB1,IL18BP,IRGM,ITGB7,PF4,PSMB8,PSMB9,SLAMF8,STAT1,TAP1,Tgtp1/Tgtp2,ZBP1 |
| Ifnar | group | 1,57×10^-18 | B2M,CASP1,CD74,CD86,GBP2,HLA-A,IFIT2,IFIT3,ISG20,Mx1,PSMB8,PSMB9,STAT1,TAP1,ZBP1 |
| ZBTB10 | transcription regulator | 3,31×10^-18 | 9930111J21Rik1/Gm12185,B2M,CD86,GBP4,GBP6,Gm5431,Ifi47,IFIT1,IFIT2,IFIT3,Igtp,Iigp1,IRGM,ISG20,Mx1,Tgtp1/Tgtp2,ZBP1 |
| Ttc39aos1 | other | 5,68×10^-18 | Apol9a/Apol9b,DDX60,GBP2,Ifi27,IFIT1,IFIT2,IFIT3,Igtp,Mx1,OAS3,PHF11,SLAMF8,TAP1 |
| PIK3CG | kinase | 8,21×10^-18 | B2M,CD74,GBP2,GBP4,GBP6,Gbp8,Gm4951,HLA-A,Igtp,Iigp1,IRGM,STAT1,TAP1,Tgtp1/Tgtp2,ZBP1 |
| ZC3H12C | other | 1,01×10^-17 | 9930111J21Rik1/Gm12185,GBP2,GBP6,Gm12250,Gm4951,Igtp,Iigp1,IRGM,SLAMF8,STAT1,Tgtp1/Tgtp2 |
| TREX1 | enzyme | 5,43×10^-17 | B2M,CASP1,CD86,DDX60,GBP4,IFIT1,IFIT2,IFIT3,ISG20,OAS3,PSMB8,PSMB9,STAT1,TAP1,TRAFD1,ZBP1 |
| STAT2 | transcription regulator | 1,70×10^-16 | CD86,CIITA,GBP6,Ifi47,IFIT1,IFIT2,IFIT3,Iigp1,Mx1,PSMB8,STAT1,ZBP1 |

Supplementary Table 4 Downregulated in IRF1 KO HSCs – Hallmark

| Term | Overlap | P-value | Adjusted P-value | Odds Ratio | Combined Score | Genes |
|---|---|---|---|---|---|---|
| **Interferon Gamma Response** | 26/200 | 2,78E-26 | 9,74E-25 | 27,2452107 | 1603,21383 | CD86;CIITA;DDX60;IFIT3;PSMB10;IFIT2;IL18BP;CASP1;ITGB7;B2M;GBP4;ZBP1;GBP6;BATF2;CD74;STAT1;MX1;TRAFD1;TAP1;PSMB8;PSMB9;ISG20;IFI27;IL7;OAS3;PSME2 |
| **Interferon Alpha Response** | 18/97 | 2,45E-21 | 4,29E-20 | 38,8657791 | 1844,52121 | BATF2;CD74;MX1;TAP1;TRAFD1;DDX60;IFIT3;PSMB8;IFIT2;PSMB9;ISG20;IFI27;IL7;CASP1;PSME2;GBP2;B2M;GBP4 |
| **Allograft Rejection** | 13/200 | 9,27E-10 | 1,08E-08 | 11,3062713 | 235,15944 | CD86;ACHE;CD74;STAT1;F2R;LY75;TAP1;PSMB10;IL7;IL12RB1;GBP2;B2M;PF4 |
| **IL-6/JAK/STAT3 Signaling** | 4/87 | 0,00279102 | 0,02442141 | 7,33382762 | 43,1327979 | STAT1;IL7;IL12RB1;PF4 |
| **Apoptosis** | 5/161 | 0,00461065 | 0,03227455 | 4,89713775 | 26,3435963 | ISG20;TOP2A;F2R;CASP1;TAP1 |

Supplementary Table 5 Downregulated KEGG in IRF1 KO HSCs

| Term | Overlap | P-value | Adjusted P-value | Odds Ratio | Combined Score | Genes |
|---|---|---|---|---|---|---|
| Proteasome | 4/46 | 2,53E-04 | 0,009941994 | 14,5230769 | 120,2955184 | PSME2;PSMB10;PSMB8;PSMB9 |
| Antigen processing and presentation | 5/78 | 1,79E-04 | 0,009941994 | 10,5091855 | 90,69240965 | CD74;CIITA;TAP1;PSME2;B2M |
| NOD-like receptor signaling pathway | 7/181 | 2,21E-04 | 0,009941994 | 6,23784958 | 52,50105435 | CASP12;STAT1;OAS3;CASP1;GBP2;GBP4;GBP3 |

Supplementary Table 6 Upregulated Hallmarks in IRF1 KO HSCs

| Term | Overlap | P-value | Adjusted P-value | Odds Ratio | Combined Score | Genes |
|---|---|---|---|---|---|---|
| **IL-6/JAK/STAT3 Signaling** | 5/87 | 8,58E-04 | 0,011428433 | 7,34272754 | 51,84545677 | SOCS3;IL1R1;CSF2RB;CD14;IL18R1 |
| **Coagulation** | 6/138 | 0,00114174 | 0,011428433 | 5,49330731 | 37,21825527 | DPP4;MMP14;SPARC;ACOX2;MMP2;CTSH |
| **TNF-alpha Signaling via NF-kB** | 7/200 | 0,00158728 | 0,011428433 | 4,39666091 | 28,33969773 | EFNA1;SOCS3;EGR1;CEBPD;NFIL3;BCL3;FOSL2 |
| **Epithelial Mesenchymal Transition** | 7/200 | 0,00158728 | 0,011428433 | 4,39666091 | 28,33969773 | EFEMP2;MMP14;SPARC;LGALS1;ITGB5;MMP2;TGFBI |
| **Inflammatory Response** | 7/200 | 0,00158728 | 0,011428433 | 4,39666091 | 28,33969773 | IFITM1;MMP14;IL18RAP;IL1R1;IL10RA;CD14;IL18R1 |
| **Hypoxia** | 6/200 | 0,00715161 | 0,036779715 | 3,72595029 | 18,40775033 | EFNA1;NFIL3;TGFBI;VLDLR;PPARGC1A;FOSL2 |
| **Xenobiotic Metabolism** | 6/200 | 0,00715161 | 0,036779715 | 3,72595029 | 18,40775033 | CYP27A1;ACOX2;IL1R1;PDK4;RAP1GAP;PTGES |

Supplemementary Table 7 HSPC gene sets

| CLP | MkP | pGM | preCFU-E | HSC_up_aged | HSC_down_aged |
|---|---|---|---|---|---|
| na | na | na | na | na | na |
| MZB1 | 1700019A02RIK | SPPL2A | CHCHD10 | SBSPON | GM3579 |
| 2010300C02RIK | 1700029M03RIK | 2610008E11RIK | ELL2 | PLSCR2 | PISD-PS3 |
| TCEANC2 | 1700063H04RIK | VSIR | APLP2 | TDRD9 | PLAC8 |
| 2900026A02RIK | 4930429B21RIK | 9130019O22RIK | CPOX | CLU | HBA-A1 |
| 3100003M19RIK | 4933402E15RIK | AAGAB | GM30289 | WWTR1 | MPO |
| 4930452B06RIK | 4933408M05RIK | ABCA9 | STX2 | OSMR | FAM46A |
| 4930539E08RIK | 4933411E08RIK | ABCD2 | 5730508B09RIK | ZG16 | HMGA2 |
| 4933402N22RIK | PROSER2 | ACOT1 | AQP9 | FAP | CAMK1D |
| 4933432K03RIK | 5830403F22RIK | ACOT7 | ARAP3 | EYA4 | MMP2 |
| 5033404E19RIK | FAM212A | ADAM9 | NADK2 | CLCA1 | BAZ1B |
| 5530400N10RIK | NOL4L | ADIPOR2 | TULP4 | NUPR1 | RGS7BP |
| 5730488B01RIK | A630038E17RIK | ADNP2 | MNS1 | DDR1 | LGALS1 |
| 6330403A02RIK | ABCA4 | AFP | RPIA | TM4SF1 | EAR-PS2 |
| SLC9A7 | ACHE | AI987944 | MFHAS1 | CCBE1 | CSF2RB |
| SSC5D | ACTN1 | AK4 | USP24 | TRPC1 | EBI3 |
| AA467197 | ACTN4 | AKR1B10 | CELA1 | GPR183 | HPGD |
| ABL1 | ADAMTSL5 | AKTIP | PEX11A | CLDN5 | HNF4A |
| ACTN3 | CD226 | ALAS1 | FZD7 | TC2N | SOCS2 |
| AGPAT9 | AK1 | ANXA2 | IL1RL1 | SELP | CD86 |
| AI467606 | ALOX5 | ANXA3 | BYSL | RIPK4 | DDX4 |
| AI661453 | ALS2CL | ARFIP1 | ATP1B2 | TMEM215 | MC5R |
| AIF1 | ANK3 | ARMC8 | SNUPN | CSPRS | P2RY14 |
| AIM1L | ANKRD29 | ARMCX2 | MINPP1 | GABRA4 | RASSF4 |
| AIRE | GLB1L | ARPC1B | ADAMTS3 | MT1 | 6030498E09RIK |
| AKAP12 | ARHGAP29 | ATG10 | FECH | CD38 | NCF4 |
| AKAP9 | ARHGAP6 | ATP6V1D | BLVRB | AGTR1A | RNASE6 |
| AL023008 | ARHGEF2 | ATXN1 | HDGF | GSTM2 | SATB1 |
| ANKRD13D | ARHGEF40 | B4GALT4 | CYB5R3 | ADGRG2 | PELI2 |
| ANKRD6 | ARRB2 | TNFSF13 | DMWD | TMEM56 | NRK |
| ANXA6 | ARSK | BCL6 | DPY19L1 | KLRB1C | EVL |
| AP3M2 | ASAP2 | BFAR | REC8 | MAF | PLXDC2 |
| APOA1 | ATP2B2 | BLOC1S2 | 1700097N02RIK | RAMP2 | SELL |
| AR | ATP2C1 | BPIFC | GORASP1 | SOCS3 | HIF1A |
| ARHGAP25 | B230217C12RIK | BTBD3 | MAPK9 | JAM2 | ITGA4 |
| ARHGEF1 | BB236558 | BZRAP1 | SSSCA1 | GM10419 | JAKMIP1 |
| ARID4A | BBIP1 | C1RA | CWF19L1 | NTF3 | IGF1 |
| ARPP21 | CCM2L | C3 | STOM | PTPRK | RBM5 |
| ATP1B1 | TTC41 | CAMK1 | DAAM1 | CLEC1A | MS4A6C |
| ATXN7L3B | BCL2L1 | CASP12 | MUL1 | GDA | ENAH |
| AURKC | BECN1 | CCBL2 | PRELID2 | GADD45G | GATA1 |
| AW061096 | BGN | CCL27A | GNA14 | NTN4 | TNFRSF13C |
| B4GALT1 | BMP2K | CCL9 | PRKAB1 | PCLO | NEDD4 |
| BAIAP2 | BMPER | CCPG1 | HOMER1 | BMPR1A | DNMT3A |
| BANK1 | BTBD9 | CCR2 | ERMAP | RARB | DSCC1 |
| BAZ2A | C1GALT1 | CD1D1 | RCL1 | PGR | MGST1 |
| BCL11A | C77649 | CD1D2 | NMT2 | MUC1 | CDCA8 |
| BFSP2 | C85328 | CD300A | RECQL | NEO1 | ATIC |
| BHLHA15 | CAP2 | CD44 | SETD6 | TMEM47 | FGF11 |
| BLK | CAPN3 | CD48 | NFRKB | CNTN1 | CREBBP |
| BMF | CASP14 | CD68 | GM12338 | EPCAM | SLC22A3 |
| BST1 | CBR3 | CDC42SE2 | ATP6V0C | SULT1A1 | ESPL1 |
| PEAK1 | CC2D2A | CEBPA | CD59A | B3GALT1 | IFNGR2 |
| C79017 | CCND3 | CEBPD | SPRY4 | CPNE8 | SGOL1 |
| C80012 | CCNJL | CHMP2B | LOC105242468 | SDPR | NCF1 |

| | | | | | |
|---|---|---|---|---|---|
| C920009B18RIK | CD180 | CLEC10A | GRWD1 | ABCA4 | KLF1 |
| CACNB1 | CD9 | CLEC4E | RNF43 | MEIS2 | RPS4L |
| CAMKV | CDC25B | CLEC7A | TFRC | ISL1 | ARHGAP30 |
| CARD11 | CDCP1 | CLOCK | DDX39 | SSPN | SMC2 |
| ACKR2 | CDK5R1 | COL4A2 | PKLR | ASPA | CAR1 |
| CCR7 | CDKN2D | CPA3 | SPHK1 | MT2 | HIST1H2AO |
| CCR9 | CEACAM1 | CPT1A | PAQR9 | SFRP1 | LRR1 |
| CD163L1 | CITED2 | CRISPLD1 | SLC38A5 | ENPP5 | CHEK1 |
| CD244 | CKAP2L | CSF1 | ZG16 | BMP4 | MS4A6B |
| CD247 | CLEC1B | CSF2RB | GPC4 | PTGFRN | RHOBTB3 |
| CD37 | CLU | CSRP2 | COL5A1 | EHD3 | SYNE2 |
| CD47 | CMAS | CTSB | MT2 | LHFP | UBTF |
| CD79A | CNNM4 | CTSC | KEL | CYP26B1 | BANK1 |
| CDC45 | CNST | CTSG | MYH10 | PERP | CDCA5 |
| CDK5R1 | CNTLN | CTSH | PRSS50 | GPX8 | MELK |
| CDV3 | CNTN1 | CTSZ | FAM132A | CYSLTR2 | 2810417H13RIK |
| CECR2 | CNTN2 | CX3CR1 | ABCG4 | VWF | TOP2A |
| CES4A | CORO2B | DHRS7 | CCDC68 | SCD1 | ECT2 |
| CHST2 | CPA4 | DNAJC28 | GML | MATN4 | HBB-B1 |
| CISH | CPD | DSTYK | COX6B2 | C4B | ITGB5 |
| CMAH | CPEB4 | DTX3L | CDR2 | PLK2 | TIMELESS |
| CNN3 | CPLX2 | EDEM1 | TSPO2 | PLCL1 | CEP350 |
| CNTROB | CRY1 | EDEM2 | 4933431E20RIK | VLDLR | PAQR8 |
| COBLL1 | CSGALNACT1 | EIF4E3 | MTHFD1 | PLEK | TM6SF1 |
| COL19A1 | CTF1 | ELANE | UBLCP1 | RND3 | EIF3J1 |
| COX6A2 | CTNNAL1 | EPHA7 | TMEM56 | ALCAM | SPC25 |
| CRIP1 | CTTN | ERO1LB | ASB17 | ACPP | PTGR1 |
| CRISPLD2 | CUEDC1 | EVI2B | ALDH1A7 | EPAS1 | KCTD14 |
| CSF1 | CYP24A1 | F13A1 | MMP14 | RGN | ARMCX4 |
| CSN3 | CYP2C29 | FCGR3 | MYO1D | EXOC3L4 | SH2D5 |
| CTLA4 | CYP4B1 | FGL2 | CENPV | ALDH1A1 | CDCA2 |
| CTNND2 | D130037M23RIK | FNDC3B | 44812 | RORB | TXNRD1 |
| CTR9 | TBC1D32 | FRS2 | ATP7B | TMEM254A /// T | LST1 |
| CTRL | D630039A03RIK | FUCA2 | PTDSS2 | CYB561 | CCNB2 |
| CYB561A3 | DAB2IP | FUT7 | HEBP1 | DENND5B | BIRC5 |
| D130062J21RIK | DACH1 | GALM | NTN4 | SEMA6D | CENPK |
| LDLRAD4 | DBH | GALNT6 | SLC11A2 | DSP | RAD54L |
| D2ERTD210E | DENND2D | GBP2 | HIF3A | ABCB1A | CCNA2 |
| D8ERTD82E | DGKG | GBP3 | PVT1 | ZSWIM5 | FAM105A |
| DBH | DLG3 | GDI2 | CASP3 | DHRS3 | MEI4 |
| DDX50 | DNER | GDPD3 | RAB4A | FAM169A | APCDD1 |
| DOC2G | DOCK1 | GLB1 | HARS | LPL | CDK5RAP1 |
| DSCAM | DPF2 | GLCE | SLC40A1 | ABAT | KIF23 |
| DTX2 | DUSP3 | GLUD1 | PLXDC1 | DSC2 | MYH10 |
| NIM1K | DZIP3 | S100A11 | NFIA | DNM3 | MAP2K7 |
| EBF1 | EHD3 | GM13152 | ACSL6 | CALML4 | RNPEP |
| EFHD1 | ELOVL7 | GM13305 | GAREM | JAG1 | IPCEF1 |
| EID2 | ENGASE | CCPG1OS | AHSP | FAM46C | COL4A2 |
| EMP1 | ESAM | GNG12 | AMPD3 | HOXB6 | PHLDB2 |
| ENDOU | ESRRB | GPI1 | RHD | PHF11C | EPB41L3 |
| ENHO | EXOC3L2 | GSDMD | TSPAN8 | MMP14 | RAD51AP1 |
| ETS1 | F10 | GUSB | PIP4K2C | TMEM30B | ANXA6 |
| ETV1 | F11R | GXYLT1 | SLC22A23 | CYYR1 | KNTC1 |
| ETV3 | F2R | H13 | MRPL52 | KLHL4 | IGF2BP2 |
| EVL | F2RL2 | HDC | AI661323 | ITGB3 | TPX2 |
| EVPL | F5 | HK3 | EIF4EBP1 | PKP2 | BTNL9 |
| FABP3 | FAM110C | HMGCS1 | ANK1 | NRG4 | AFF3 |

| | | | | | |
|---|---|---|---|---|---|
| FADS3 | MVB12B | HOMEZ | OAF | PFN2 | CDK1 |
| FAM129C | ZC2HC1B | HP | FAM213A | XDH | COTL1 |
| FAM155A | FAM171A2 | HPS3 | IGSF3 | CLEC7A | CENPN |
| FAM167A | FAM64A | IBTK | EPHX2 | EFEMP1 | SNCA |
| FCRL1 | FEM1A | IDH1 | REC114 | EXOC3L2 | COL4A1 |
| FGF4 | FERMT3 | IFT46 | NXPE2 | A730089K16RIK | UHRF1 |
| FOXO1 | FHL1 | IGSF6 | AQP11 | ID2 | AI451250 |
| GBP8 | FKBP1B | IKZF2 | DCAF4 | GEM | LCORL |
| GFRA1 | FLI1 | IL15 | HPN | OCLN | SORL1 |
| GFRA2 | FNBP1L | FAM234A | TNFAIP2 | IRF2BPL | 4930515G01RIK |
| GGTA1 | FOXD2 | ITGB7 | FAM234B | KDF1 | TCF19 |
| GIMAP4 | FSTL1 | KANSL1L | REPS2 | CTTNBP2NL | BUB1 |
| GIMAP5 | FYB | KCTD12B | PTPN14 | MAB21L2 | DNMT3B |
| GIMAP7 | GADD45G | KCTD20 | AP1B1 | FFAR2 | AP1S2 |
| GIMAP9 | GAS2L3 | KLHL5 | OPTN | GPRIN2 | DEPDC1B |
| GM16340 | GATA2 | LAIR1 | GM3417 | NXNL2 | GM14085 |
| GM16364 | GCHFR | LAMP2 | CDS2 | AR | CDKN1B |
| GM19576 | GDA | LDHB | SNHG3 | KDR | TET1 |
| TSC22D1 | GHR | LGALS1 | LMNA | CHST11 | CKS2 |
| PAX5 | GLRP1 | LMO4 | CLDN13 | EPDR1 | DHFR |
| ARIH1 | LOC545966 | LTA4H | 2210016F16RIK | KIF21A | USP37 |
| GM3579 | RBPMS2 | LY6C1 | USP6NL | PTGER4 | PHLDA2 |
| GM5914 | GM9798 | MANBA | APLF | MYOM1 | CORO2B |
| GPR174 | GMPR | MAPKAPK2 | RHAG | AMPD3 | CNTRL |
| GPR25 | GNAZ | MAPKAPK3 | MTHFD1L | NUDT10 | LTN1 |
| A330009N23RIK | GNB5 | MCPT8 | RMDN3 | FHDC1 | BRCA1 |
| GTPBP2 | GNG11 | MET | ST8SIA6 | ENKUR | CD34 |
| HAAO | GP1BA | MGST1 | SPIRE1 | RASSF10 | KCNK12 |
| MROH2A | GP1BB | MORC3 | AMIGO2 | CLEC1B | TACC3 |
| HIP1R | GP1BB | MRGPRA2A | CCNE1 | PDE9A | MCM5 |
| HIVEP2 | GP9 | MS4A3 | MBOAT2 | DOCK9 | AI504432 |
| HMG20A | GPR150 | MS4A6B | ZMAT3 | ALDH5A1 | NOCT |
| HSD17B13 | ADGRG2 | MS4A6C | PLXNA1 | RNF150 | 1700023H06RIK |
| ICOS | GRTP1 | MAP4 | COX6A1 | CEBPD | PRIM1 |
| IGH-VJ558 | GSTT1 | MAP7 | CACHD1 | RHOJ | RRM2 |
| IGHD | GSTT3 | MTUS1 | RHOF | CD200R4 | ERCC6L |
| IGLL1 | GUCY1A3 | MYO5A | PITRM1 | PCDHB16 | CDT1 |
| IL10RA | GUCY1B3 | NCEH1 | IFNGR1 | RAB40B | MCM10 |
| IL13RA2 | GULP1 | NCOA4 | KIF26B | SGK1 | 1700097N02RIK |
| IL17RC | H1FX | NCOA7 | ABCB7 | SNHG14 | TYMS |
| IL7R | HBEGF | NPC2 | EPS15 | SEMA7A | BC035044 |
| IMPA1 | HDGFRP2 | NPL | MYLPF | C130026I21RIK | PDGFC |
| INADL | HDGFRP3 | NRP1 | PTK2 | GM5148 | LHFPL2 |
| INCA1 | HDLBP | NRXN1 | ZFP800 | RASEF | AK4 |
| IRF2 | HIF3A | NSMAF | 1700006J14RIK | TRIM47 | ANXA2 |
| ITPR2 | HIPK2 | OASL2 | CPSF2 | NXPE2 | FLT3 |
| ITSN1 | HNF4A | PAPSS2 | PHB2 | SLC7A7 | SYCE2 |
| IZUMO4 | HOMER2 | PARP16 | DNAJA3 | CTSC | CD37 |
| KCTD1 | ICA1 | PARVG | SPTA1 | ETL4 | BPTF |
| KIDINS220 | IFITM3 | PELI2 | LRWD1 | BHLHE40 | UNG |
| KIF23 | IGF2BP2 | PGAM1 | LRRC20 | RDH10 | CLNK |
| KIFC2 | IGSF21 | PHLDB2 | MPV17L2 | FHL1 | SKP2 |
| KLF3 | ITGA2B | PROCR | SAMM50 | PHACTR1 | MAMDC2 |
| KLHL30 | ITGA8 | PRTN3 | GSTP1 | STXBP4 | BUB1B |
| KMO | ITGB3 | PSMB8 | MDH2 | 1810073O08RIK | SYNPO2 |
| KRT24 | ITM2A | RAB11FIP5 | PSMC2 | GHR | CKS1B |
| LAT | JPH1 | RAB3D | NAT6 | GDPD1 | GIMAP7 |

| | | | | | |
|---|---|---|---|---|---|
| LAX1 | KALRN | RAB44 | B3GALNT2 | MYOF | KIF18B |
| LCK | KCNC1 | RASSF4 | PKN2 | RNF128 | FILIP1L |
| LIFR | KCNJ5 | RASSF8 | ACOT6 | CASP12 | SPSB4 |
| LPAR1 | KDR | RILPL2 | CARNMT1 | INTS6 | LIG1 |
| LRP1 | KRTAP15 | RNF149 | MC2R | LRRN1 | CKAP2 |
| LY6D | P3H2 | RNPEP | LRIG1 | SYNM | MTHFD2 |
| LYNX1 | P3H3 | ROPN1L | TSPAN33 | FAT1 | SYK |
| MALAT1 | LHFP | RORA | NECAP2 | LDHD | MBP |
| MAMDC4 | LHX3 | RPL7L1 | PCMT1 | CPEB2 | CTR9 |
| MAP2K7 | LIMS1 | RRM2B | SCARB1 | OXR1 | ENDOD1 |
| MAP4K2 | ADGRL1 | RTN3 | GEMIN5 | AMOTL2 | NCAPH |
| MAPT | LRPAP1 | RUFY1 | AI854703 | TBC1D8 | POLE2 |
| MARCKS | LRRFIP2 | SCCPDH | RYK | TOX | KPNA4 |
| MCTS2 | MAGED2 | SCPEP1 | STEAP3 | ARHGAP29 | ASF1B |
| MESP1 | MANSC1 | SEPP1 | PABPC4 | TACSTD2 | SKA1 |
| MID1 | MDC1 | SHMT1 | RCC2 | GIPC2 | LMNB1 |
| MMP11 | MDM1 | SIKE1 | DAPK2 | CTSW | CKAP2L |
| MRGPRE | MED12L | SLC25A12 | FPGS | SPO11 | RFC2 |
| MSI2 | MFAP2 | SLC35B4 | PIGQ | PRKD1 | CNR2 |
| MTA3 | MFAP3L | SLC35C2 | ATP4A | PCDHB17 | SACS |
| MYO1C | MMRN1 | SLCO3A1 | EMC9 | 4930550C14RIK | MLEC |
| N4BP3 | MPDZ | SLFN9 | AA408650 | LPAR6 | ERG |
| NEFH | MPL | SMARCAD1 | PM20D2 | PROS1 | TMSB10 |
| SAPCD1 | MRVI1 | SMEK2 | PLA2G4C | GPR4 | SLC29A3 |
| NKX2-6 | MAP1A | SP100 | SLC19A1 | ST3GAL6 | MB21D1 |
| OBFC1 | MYL3 | SPG21 | SRM | DZIP1 | STIL |
| OIT1 | MYOM1 | SPSB4 | SLC26A1 | ARHGEF28 | CIT |
| P2RX3 | NBEA | SSBP1 | ST3GAL5 | ABCB1B | SLC16A7 |
| P2RY10 | NCOA3 | ST3GAL2 | NOP2 | CCNJ | VIM |
| P2RY13 | NDN | STXBP4 | SPPL2B | ROR2 | GINS1 |
| PARD6G | NDST3 | STXBP6 | API5 | ZFP503 | ORC1 |
| PARP3 | NSMF | SWAP70 | UBXN2B | NCKAP1 | ANKRD29 |
| PDE1B | NFS1 | TAPBPL | URGCP | H2-EB1 | DLGAP5 |
| PDE2A | NICN1 | TBC1D9B | KCNG2 | ASB4 | CEACAM1 |
| PDZD4 | NRARP | TCF7 | SLC41A3 | ART4 | MARCKSL1 |
| PFKL | NRGN | TCP11L2 | NUP85 | PIM2 | H2AFX |
| PHF2 | NSMCE1 | LOC665506 | PPM1G | ZFP932 | CFP |
| PIK3R1 | OBSL1 | TCRG-C4 | TMEM238 | HOXB3 | ARHGAP11A |
| PIK3R6 | OPHN1 | TFEC | ACSL1 | PBX3 | NMRAL1 |
| PIP4K2A | OPLAH | TIPARP | CCT3 | CD55 | GMNN |
| PLEKHA7 | OPRM1 | TMCC2 | ADD2 | BCL6 | CENPM |
| PLEKHG2 | P2RX1 | TOB1 | BC021614 | PDGFD | C1QC |
| POGZ | PARD3B | TOR3A | TXNRD2 | SDC4 | HMGA1 |
| POLR1A | PBX1 | TREM3 | DTL | NEAT1 | ABLIM1 |
| POU2AF1 | PCCA | TRIM30D | CSNK2A1 | PRCP | HN1 |
| PPP3CA | PCYT1B | TSPAN3 | VKORC1L1 | SAT1 | WDHD1 |
| PPP3CC | PDCD4 | TTC39C | SCLY | FARP1 | ANKHD1 |
| PRKCB | PDE10A | TXNDC11 | STRA13 | VMP1 | EZH2 |
| PRSS53 | PDE5A | UAP1L1 | TBCCD1 | ARPIN | CLSPN |
| PTEN | PDLIM7 | UBE2B | 9530077C05RIK | IL18R1 | SDF2L1 |
| PUF60 | PDZK1IP1 | UBE2R2 | ATG4B | RGMB | PIK3AP1 |
| PYDC4 | PEAR1 | USP3 | ARHGEF25 | DHX40 | AKR1C12 |
| PYGM | PF4 | VLDLR | USP47 | NPDC1 | KCNAB2 |
| RABEP2 | PHLDA1 | VRK2 | ATP13A3 | ERP27 | GM9706 |
| RAG1 | PHLDB1 | XRCC4 | FANCA | 2310001H17RIK | ATAD2 |
| RALGPS1 | PIGN | ZC3H7A | POLRMT | CELF5 | HIST1H4I |
| RASAL1 | PIGZ | ZC3HC1 | MTX1 | IL1RAPL2 | CD5L |

| | | | | | |
|---|---|---|---|---|---|
| RASGRP3 | GSAP | ZFP184 | INO80E | HOXB5 | NRM |
| RBM47 | PIP5K1B | ZFP383 | POLR2L | ENPP1 | RRM1 |
| RBP7 | PITPNM1 | ZFP760 | NOL10 | AK1 | GM16340 |
| RCSD1 | PKD2 | ZSCAN21 | HACD3 | NXPE4 | ARRB2 |
| RDH12 | PKP3 | | KLHL22 | GREB1 | CEP85 |
| RGS7BP | PLAGL1 | | PAQR4 | KCNK6 | 1700019G17RIK |
| RGS8 | PLCB1 | | SCRN3 | AKR1C19 | POLD1 |
| RILPL1 | PLEK | | DHODH | ETS1 | FAM167A |
| RPGRIP1 | PLS1 | | FAM134A | PIK3R1 | VRK1 |
| SATB1 | PLPP3 | | RPS6KA4 | MFAP3L | MCM3 |
| SCN4B | PPBP | | ELAC2 | SLC6A15 | CDC20 |
| SDC1 | PPFIA3 | | ADD1 | EPHX1 | DYNLL1 |
| SDC4 | PPFIBP1 | | NOL9 | PRTN3 | FAM64A |
| SEMA4A | PPIC | | SORBS1 | SERPINB6A | SMC1A |
| SEMA4D | PRAF2 | | PRICKLE4 | CCDC160 | RBM26 |
| SERF2 | PRAME | | GLG1 | NAV1 | PHGDH |
| SGSH | PRKCA | | CS | C530008M17RIK | EMID1 |
| SGSM1 | PRKCDBP | | UBQLN4 | S100A6 | PRIM2 |
| SLA2 | PRKCQ | | NUP133 | NOVA1 | RIN3 |
| SLC17A9 | PRKD1 | | NDST2 | THBD | RPRD1B |
| SLC18A1 | PRKG1 | | SLC35E2 | EVC2 | ISLR |
| SLC22A6 | PROS1 | | GPAM | FADS3 | FAM78A |
| SLC22A8 | PRR11 | | TMEM41B | GPLD1 | OAS2 |
| SLC37A2 | PRSS2 | | RABGEF1 | ST3GAL1 | MAP10 |
| SMAD7 | PSD3 | | AKR7A5 | YPEL2 | UMPS |
| SORCS2 | PSEN1 | | ANAPC15 | TRIM2 | FEN1 |
| SOST | PSTPIP2 | | CDYL | TGFB3 | TRIP13 |
| SPAG8 | PTGER3 | | IBA57 | TGM2 | SHMT1 |
| SPRY2 | PTGS1 | | KCTD7 | SH3BGRL2 | IGFBP4 |
| SRPK3 | PTN | | TBL2 | NR3C2 | DHRS7 |
| SRSF3 | PTPRF | | ZDHHC14 | SEMA3E | DNA2 |
| ST3GAL6 | PTRF | | MCCC1 | SLAMF1 | CDC25A |
| ST6GAL1 | PTTG1IP | | IGTP | EVC | REEP5 |
| ST8SIA1 | PYROXD2 | | GLS2 | AJUBA | E2F2 |
| STAM2 | RAB11A | | FAM136A | FADS6 | CMC2 |
| STAMBPL1 | RAB27B | | SLC35D1 | ITGA6 | DDX39 |
| STC1 | RAB37 | | DAZAP1 | PYGM | NUP85 |
| SULF1 | RAB3D | | FBXO30 | CYTIP | FRAT2 |
| SYNJ2 | RABGAP1L | | KLK1 | SHTN1 | TIGD3 |
| SYNPO2L | RALGPS2 | | CCBL1 | ZC3H12D | HMMR |
| SYTL2 | RAP1B | | GLCCI1 | PIP5K1B | CABLES1 |
| TBC1D16 | RAP2B | | SPECC1 | SEMA4F | SRM |
| TBXA2R | RAPGEFL1 | | 1600002K03RIK | RPS6KA3 | FCER1G |
| TECPR1 | RAPSN | | RECQL4 | EXOC6B | NUTF2 |
| TLR7 | RBMS2 | | SLC7A5 | FBXO2 | TRIP12 |
| TMED2 | RGS1 | | PPP2R1B | PPP1R16B | FCHO1 |
| TMEM108 | RGS18 | | MRPS36 | WWP2 | NFAM1 |
| TMEM119 | RHOBTB3 | | AARSD1 | PPIL6 | DOCK8 |
| TMEM121 | RHOF | | ATAD3A | SPON1 | PGP |
| TMTC1 | RNF180 | | CCNC | TMEM163 | AI746446 |
| TOPBP1 | ROBO3 | | INTS5 | NREP | PMF1 |
| TOX3 | RXRA | | PDE12 | MYO6 | FKBP2 |
| TPM2 | SARDH | | GJA1 | SELM | MCM7 |
| TREM1 | SCARF1 | | TSTD3 | FZD1 | HAUS8 |
| TRIB3 | SCRT2 | | AHCTF1 | DPY19L2 | MCM2 |
| TRIM26 | SDPR | | NDUFA12 | EFNA1 | ST3GAL5 |
| TRP53I11 | SEMA6D | | EPDR1 | TRPM6 | ATP13A2 |

TRP53INP1
TSNAXIP1
UACA
URB1
VAMP4
VPREB1
VPREB1
WNK2
XRCC6
NDUFA13
YPEL1
ZAP70
ZBED6
ZBTB2
ZBTB20
ZFP503
ZFP629
ZMIZ1

SERPINA12
SERPINA1B
SERPINE2
SGCD
SGCE
SHROOM4
SIK3
SLA
SLAMF1
SLC14A1
SLC14A2
SLC16A2
SLC25A44
SLC38A4
SLC45A2
SLC6A3
SLC6A8
SMO
SOD1
SORBS3
SPTB
SPRR2H
SRGAP3
STAM2
STAU2
STIM1
TAGLN2
TBC1D4
TBXAS1
TEK
TGM2
THBD
TIE1
TIMP3
TMEM158
TMEM174
TMEM43
TMEM59
TMEM98
TMIE
TNFAIP8L1
TNIK
TPX2
TRBV13-2
TREML1
TRIM47
TRPC6
TUBA8
TULP3
TYRP1
UBASH3B
UCP1
UFSP1
UNC119B
USH2A
USP40
USP54

DNAJC30
STK24
NAT10
MYDGF
RAB26OS
NOP10
LARS2
NXPE4
REXO4
ABCA3
SFXN1
ARRB1
NVL
MTFR1L
CHST1
JMY
FGFRL1
AP2B1
DHRS11
EPT1
MTR
ECD
BC017158
HEATR1
RFFL
COQ4
LRRC8C
RAB22A
ASPH
ZFP598
VARS2
TXNRD3
TOM1
ADCK5
CPEB3
GPATCH1
TRIM37
SREBF1
MPG
LOC102641161
BTAF1
SLC2A8
FARSB
MIF4GD
WDR18
DNAJA4
AAAS
LMCD1
PSMD11
METTL13
TMEM209
CMC2
URB1
NAA40
XPO4
CARM1
DPF3

APOL7B
TNFAIP2
BBOF1
SERPINB8
GCA
FZD3
AQP9
RBPMS2
FAM135A
ADCK3
2610305D13RIK
PAWR
TXNIP
CAMKK1
SRD5A1
RAB34
MYLK
EZH1
H2-AA
TMEM204
COL16A1
FAM84B
TMTC2
KRT8
PHKA1
RORC
WTIP
CDKL2
RASSF6
MEIOB
PAQR7
C1RA
ZFP462
ADCY9
RPRD2
CREBL2
MTMR7
SBF2
TSPAN17
TNFSF10
ARHGAP6
FERMT2
2010107G23RIK
CPM
LOXL3
GPR146
MYO5C
SHROOM3
UNC13B
CHRNB1
ARAP2
NDRG1
LOX
ZFAND5
PRNP
SNX24
GM8267

H2AFV
SETD8
GMEB1
NEFH
RAD23A
SOX4
FANCI
CYCS
PES1
MCM4
NSG1
GM6756
ARHGDIB
PBX2
AAAS
WDFY2
NOP56
BC147527
TNFRSF18
MCM6
NUP155
DONSON
NCL
DTYMK
SCARB1
SNRPA
WEE1
PA2G4
ELOVL5
NUDC
NUP62
E130309D02RIK
MEAF6
MEX3A
PPP1R9A
COX6A1
SH3KBP1
NDUFA10
CDK2AP1
PKMYT1
CNOT11
MTX1
CACNA2D1
SELPLG
PLIN2
AP1M1
NCKAP1L
FAAH
DAZAP1
GIT2
H2AFZ
LOC102641161
CUTA
TRIM27

VASH1
VCL
VEZF1
VPS53
VWF
WDR37
NPM2
ZEB1
ZEB2
ZFP111
ZFP169
ZFP36L1
ZFP385A
ZFP410
ZFP697
ZFP948
ZYX

EXOSC4
ORC2
UNC5CL
BC055324
DAGLB
USP19
RBM14
POT1A
SSBP3
ZFP652
RGS9BP
RRBP1
IARS
PSMB3
FNIP2
C1QBP
TIMM44
TIMM10B
MRPS2
UGCG
SCFD2
ZZEF1
CERCAM
ZFP142
PPP1R11
HSPH1
PA2G4
NUP155
PLA2G12A
GNL2
MTAP
DAPK1
TRMT6
TNKS
USP31
LIG3
PPP2R4
E130309D02RIK
GAR1
CCSAP
SERPINB9B
SPRYD3
OLFM1
RAD23A
PRKAR2A
CLIC4
BC051142
ARFGAP1
INPP5F
HSF1
LSS
CCDC186
RCOR3
PDAP1
CNNM2
DCAF7
TXN2

IRF6
FAM63A
PAK6
TMEM158
HOMER1
BCL3
CAMK2N1
TMEM117
ZFP334
ADCY6
FYB
A630072M18RIK
FAM184A
E2F5
RBM19
RIN2
A630033H20RIK
CBR3
SLC35F2
TROVE2
IL6ST
LONRF2
ABCA5
ISCA1
CARD10
NABP1
CXCL16
PBX1
RRAGD
TRIB3
DNAL1
SERPING1
PPM1E
SPATS2
LGMN
ATP10A
MYO1E
MEF2C
ZFP950
MYO1D
MYCT1
CLEC14A
PROCR
MBOAT2
GALNT6
PLSCR4
RPRM
SORBS1
GKN3
EPS8
ARHGAP5
LANCL3
B4GALNT4
ALDH3A1
2310030G06RIK
NACC2
4833418N02RIK

| | |
|---|---|
| SMPD4 | ROCK2 |
| ABCF1 | ACSL4 |
| CAPRIN2 | CLIC5 |
| JMJD6 | ZFP612 |
| FBXW2 | KCND3 |
| JRK | H2-AB1 |
| PTPN13 | ASPH |
| TAF10 | SPINT1 |
| ESCO2 | CDCP1 |
| GM42151 | ACKR1 |
| ZFP39 | NDN |
| XKR5 | KISS1R |
| 2810021J22RIK | MIER1 |
| GATAD2A | CPEB4 |
| BEND3 | IL22RA2 |
| ENPP1 | ADGRL4 |
| WDR60 | PJA2 |
| XRCC3 | CDC14A |
| OGDH | SLC2A13 |
| METTL1 | SYNPO |
| PLVAP | PRKCZ |
| WDR43 | SMPDL3B |
| PKP2 | DCBLD2 |
| DCAF12L1 | MED12L |
| NDUFA10 | PRKAG2 |
| LOC105242484 | CYP4B1 |
| DDX47 | SRXN1 |
| TLCD1 | SMAGP |
| PPRC1 | CACNA1B |
| PAPD7 | CITED2 |
| MON1B | PPP1R12A |
| ALYREF | ENTPD2 |
| UBR5 | ERO1L |
| AGRN | GCLM |
| POM121 | TRPM4 |
| SLC44A1 | HECTD2 |
| DSN1 | FAR2 |
| IPMK | PTGER3 |
| RBM28 | PABPC4L |
| 2310061I04RIK | GSTT1 |
| BRAT1 | RCVRN |
| FANCD2 | PPP1R3B |
| GPR155 | LAMP2 |
| HNRNPU | CDC73 |
| AAR2 | 9130008F23RIK |
| TMEM131 | AA986860 |
| AGAP1 | PROM2 |
| PSMD3 | RORA |
| ARL4A | MMRN1 |
| RNF126 | ACER2 |
| 2500002B13RIK | LMBR1 |
| TTF2 | SNHG18 |
| MIR3069 | SMTNL1 |
| DPP8 | KRT18 |
| GM17399 | CRIM1 |
| DHTKD1 | WDR17 |
| ITFG2 | CLEC12B |

RNPS1
GSPT1
ORC4
44806
RFNG
CDK5RAP1
RANGAP1
TMED9
HECTD1
GNE
TBC1D25
U2SURP
CREBL2
CD81
LEFTY1
MIR1949
PRNP
PVRL2
FAM46C
FRAT1
DUS2
NKX1-2
CYYR1
HGH1
FBXO10
SLC29A2
GTSE1
IRG1
F8
MC1R
PFN4
HDAC7
TFE3
AGTR1A
ELMO3
PTPN13
PCDHB12
FGFR3
LSR
CYP11A1
CPT1C
SEC14L1
EMP2
SBSN
PCDHB20
RASGEF1B
SHROOM2
SLCO2A1
ADAM9
CRACR2B
SULF2
CXADR
TMEM38B
RCAN2
DIXDC1
PCYT1A
LPAR4
GYLTL1B
UNC45B
9630013D21RIK
EFHD1
GP1BB
CNBD2

**Supplementary Table 8: Key Resources Table**

| REAGENT or RESOURCE | SOURCE | IDENTIFIER |
|---|---|---|
| **Antibodies** | | |
| Biotin anti-mouse/human CD45R/B220 (RA3-6B2) | BioLegend | Cat: 103204 |
| Biotin anti-mouse CD4 (GK1.5) | BioLegend | Cat: 100404 |
| Biotin anti-mouse CD8a (53-6.7) | BioLegend | Cat: 100704 |
| Biotin anti-mouse/human CD11b (M1/70) | BioLegend | Cat: 101204 |
| Biotin anti-mouse Ly-6G/Ly6C (Gr-1) (RB6-8C5) | BioLegend | Cat: 108404 |
| Biotin anti-mouse TER-119/Erythroid Cells (TER-119) | BioLegend | Cat: 116204 |
| PE/Cy5 anti-mouse/human CD45R/B220 (RA3-6B2) | BioLegend | Cat: 103209 |
| PE/Cy5 anti-mouse/human CD11b (M1/70) | BioLegend | Cat: 101209 |
| PE/Cy5 anti-mouse TER-119/Erythroid Cells (TER-119) | BioLegend | Cat: 116209 |
| APC/Cy7 anti-mouse CD4 (RM4-5) | BioLegend | Cat: 100526 |
| PE/Cy5 anti-mouse CD8a (53-6.7) | BioLegend | Cat: 100709 |
| APC anti-mouse human CD11b (M1/70) | BioLegend | Cat: 101212 |
| PE/Cy7 anti-mouse CD19 (6D5) | BioLegend | Cat: 115520 |
| PE anti-mouse CD45.1 (A20) | BioLegend | Cat: 110708 |
| FITC anti-mouse CD45.2 (104) | BioLegend | Cat: 109806 |
| APC-eFluor780 anti-mouse CD117 (c-Kit) (2B8) | Invitrogen | Ref: 47-1171-82 |
| Pacific Blue anti-mouse Ly-6A/E (Sca-1) (E13-16.7) | BioLegend | Cat: 122520 |
| NK.1. Brilliant Violet 421 | BioLegend | Cat: 108722 |
| Alexa Fluor 700 anti-mouse CD48 (HM48-1) | BioLegend | Cat: 103426 |
| PE/Cy7 anti-mouse CD48 (HM48-1) | BioLegend | Cat: 103424 |
| APC anti-mouse CD150 (SLAM) (TC15-12F12.2) | BioLegend | Cat: 115910 |
| PE anti-mouse CD135 (FLT3) (A2F10)) | BioLegend | Cat: 135306 |
| PE/Cy7 anti-mouse CD105 (MJ7/18) | BioLegend | Cat: 120410 |
| PerCP-eFluor710 anti-mouse CD41 (eBioMWReg30) | Invitrogen | Ref: 46-0411-82 |
| PE anti-mouse CD41 (MWReg30) | BioLegend | Cat: 133906 |
| Alexa Fluor 700 anti-mouse CD16/32 (93) | Invitrogen | Ref: 56-0161-82 |
| FITC anti-mouse Ki67 | BD Biosciences | Cat: 556026 |
| PE anti-mouse Ki67 | BD Biosciences | Cat: 556027 |
| Brilliant Violet 510 Streptavidin | BioLegend | Cat: 405234 |
| FITC anti-mouse MHC II | Miltenyi Biotec | 130-123-666 |
| Multi ubiquitin mAb (D071-3) | MBL International Corporation | D071-3 |
| Alexa Fluor 488 goat anti-mouse IgG (H+L) Cross-Adsorbed ReadyProbes$^{TM}$ Secondary Antibody | Invitrogen | Cat: R37120 |
| **Chemicals, peptides, and recombinant proteins** | | |
| 7-Aminoactinomycin D (7-AAD) | Invitrogen | Cat: A1310 |
| Propidium Iodide | Molecular Probes | Cat: P3566 |
| Heparin solution | Stem Cell Technologies | Cat: 07980 |
| Lipopolysaccharide (LPS) | Sigma Aldrich | Product No: L4005 |
| **Critical commercial assays** | | |
| BD cytofix/cytoperm fixation/permeabilization solution kit | BD Bioscience | Cat: BD 554714 |
| BD FITC Mouse Anti-Ki-67 Set | BD Biosciences | Cat: 556026 |

| | | |
|---|---|---|
| SMART-Seq® v4 Ultra® Low Input RNA Kit | Takara Bio USA, Inc. | Cat. Nos. 634888, 634889, 634890, 634891, 634892, 634893, 634894 (091817) |
| Single Cell RNA Purification Kit | Norgen Biotek Corp. | Cat: 52800 |
| Illumina Nextera XT kit | Illumina | 4456740 |
| BD Annexin V: FITC Apoptosis Detection Kit I | BD Pharmingen$^{TM}$ | BD 556547 |
| **Deposited data** | | |
| RNA-seq data | This manuscript | [redacted] |
| **Publically available data** | | |
| Expression profiling by array | PMID: 18838472,<br>PMID: 20522712 | GSE6891 |
| **Experimental models: Organisms/strains** | | |
| Mouse: C57Bl/6JRj | In-house breeding | |
| Mouse: B6.SJL (B6.SJL-*Ptprc*$^{a}$*Pepc*$^{b}$/BoyCrCrl) | In-house breeding | Charles River, Strain Code: 564 |
| Mouse: IRF1 KO (B6.129S2-*Irf1*$^{tm1Mak}$/J) | In-house breeding | Jackson Laboratory, stock number 002762 |
| **Software and algorithms** | | |
| FlowJo | BD, Treestar | v. 10 |
| GraphPad Prism | Dotmatics | v. 9 |
| Gene Set Enrichment Analysis (GSEA) software | Broad institute | http://software.broadinstitute.org/gsea/index.jsp |
| Enrichr | Chen, E.Y., et al. (2013).<br>DOI: 10.1186/1471-2105-14-128<br><br>Kuleshov, M.V. et al. (2016).<br>DOI: 10.1093/nar/gkw377<br><br>Xie, Z. et al. (2021).<br>https://doi.org/10.1002/cpz1.90 | https://maayanlab.cloud/Enrichr/ |
| RNA Galaxy workbench 2.0 | RNA Bioinformatics Center (RBC) | https://usegalaxy.eu/ |
| TapeStation Analysis Software 3.2 | Agilent Technologies, Inc. | |

| | | |
|---|---|---|
| Cell Radar | https://karlssong.github.io/cellradar/ | https://karlssong.github.io/cellradar/ |
| **Other** | | |
| BD Aria III | Becton Dickinson | |
| BC CyAn ADP | Beckman Coulter | |
| BD LSR X-20 | Becton Dickinson | |
| NovaSeq S4 | Illumina | |
